# HOPS disruption impairs APP trafficking and processing, promoting exosomal secretion of APP-CTFs

**DOI:** 10.1101/2025.09.09.675103

**Authors:** Derk Draper, Anna E. George, Tineke Veenendaal, Suzanne van Dijk, Frederik J. Verweij, Judith Klumperman, Ginny G. Farías

## Abstract

Amyloid precursor protein (APP) is a key player in various neuronal functions but also the source for toxic Aβ that accumulates in the brain of Alzheimer patients. APP trafficking and processing depend on the endo-lysosomal system, but the molecular mechanisms that coordinate these processes remain unclear. Here, we studied the HOPS complex, a central regulator of endo-lysosomal maturation. We show that HOPS disruption impairs retromer-mediated recycling of APP to the TGN, resulting in the accumulation of APP in late endosomes. In neurons, this accumulation is spatially restricted to somatodendritic endosomes. These APP-containing endosomes are catalytically inactive and lack the γ-secretase subunit PSEN2. However, they do contain BACE1, which leads to the build-up of toxic APP C-terminal fragments (APP-CTFs) upon HOPS disruption. Notably, loss of HOPS enhances secretion of APP-CTFs by exosomes, suggesting a potential mechanism for disease propagation. Together, our findings establish a mechanistic link between HOPS dysfunction and aberrant APP processing, with implications for neurodegeneration.

**Highlights:**

- HOPS KO impairs retromer-mediated APP recycling to the TGN
- HOPS disruption redistributes APP to somatodendritic stationary late endosomes
- HOPS depletion increases APP and BACE1 convergence, causing APP-CTF accumulation
- These APP-CTFs accumulate in catalytically inactive endosomes that lack PSEN2
- APP-CTFs are secreted via exosomes, potentially promoting disease propagation

## Introduction

Alzheimer’s Disease (AD) is a major global health burden and the most common cause of dementia, leading to progressive cognitive decline. This decline is associated with synapse loss and neuronal death, which are shared features of additional neurodegenerative diseases including Parkinson’s Disease and Huntington (Wilson et al., 2023). A key driver in AD pathology is the transmembrane protein Amyloid Precursor Protein (APP), which is processed by secretases into fragments of varying properties (Müller et al., 2017). Disease-modifying mutations in APP or the secretase presenilin-1/2 (PSEN) lead to alterations in APP processing and consequently extracellular deposition of neurotoxic amyloid-β (Aβ) oligomers (Goate et al., 1991; Rogaev et al., 1995; Sherrington et al., 1995; Jonsson et al., 2012). Additionally, APP-derived C-terminal fragments (APP-CTFs) accumulate intracellularly and contribute to synaptic dysfunction and cholesterol imbalance (García-Ayllón et al., 2017; Bretou et al., 2024; Luo et al., 2025). Although genetic risk factors have been identified for familial AD, most AD cases (∼95%) have no clear genetic background and our understanding of disease modifiers that regulate APP trafficking and processing in these patients remains limited (Van Acker et al., 2019).

The endo-lysosomal system comprises an intricate membrane network of compartments that together are essential for endocytic and autophagic protein trafficking and clearance. Interestingly, defects in the endo-lysosomal system can precede clinical AD manifestations (Cataldo et al., 2000). These defects include the enlargement of endosomes and accumulation of proteolytically deficient lysosomes, which indicates a link between disrupted endo-lysosomal homeostasis and AD (Cataldo et al., 2000; Gowrishankar et al., 2015a; Kwart et al., 2019). Notably, amyloidogenic APP processing — resulting in APP-CTFs and Aβ — occurs at least partially within the endo-lysosomal system (Park et al., 2022). Of note, APP cleavage products can disrupt lysosomal function, while - vice versa - altering lysosomal activity can affect APP processing (Kwart et al., 2019; Bretou et al., 2024; Gallwitz et al., 2024; Mueller-Steiner et al., 2006). Though these findings highlight the importance of endo-lysosomal homeostasis on APP processing and clearance, the precise role of the endo-lysosomal system herein remains unclear.

The HOPS complex is a master regulator of lysosome biogenesis and functioning (van der Beek and Klumperman, 2025). This cytoplasmic complex, composed of six subunits (VPS11, VPS16, VPS18, VPS33a, VPS39, and VPS41), regulates fusion of lysosomes with late endosomes, autophagosomes, and trans-Golgi network (TGN)-derived biosynthetic carriers transporting lysosomal membrane proteins (Schleinitz et al., 2023; Sanzà et al., 2025). HOPS disruption also affects endosomal maturation, leading to enlarged, early-to-late hybrid endosomes that fail to recycle mannose 6-phosphate receptor (M6PR), which targets soluble acid hydrolases from TGN to early endosomes (van der Beek et al., 2024; Walia et al., 2024). Mutations in HOPS subunits lead to severe neurodegenerative diseases (van der Beek et al., 2019; Steel et al., 2020; van der Welle et al., 2021). For example, we and others found that patients with mutations in the *VPS41* gene display severe neuronal and cellular defects, including ataxia, dystonia and reduced autophagic flux (van der Welle et al., 2021; Steel et al., 2020). Interestingly, VPS18 and VPS41 expression levels are generally reduced in AD (Dharshini et al., 2019; Polanco et al., 2022), while overexpression of HOPS and its interactor ARL8B rescues Aβ-induced neurotoxicity in a *C. Elegans* AD model (Griffin et al., 2018). It remains unclear, however, whether defects in HOPS affect APP trafficking and processing.

Here, we investigated the effect of HOPS impairment on APP trafficking and processing in human HeLa cells and primary rat hippocampal neurons. In both models, we found that HOPS disruption led to the accumulation of APP in late endosomes and a concomitant depletion from the TGN. In neurons, this endosomal accumulation was restricted to the somatodendritic domain. Mechanistically we found that in the absence of HOPS, APP failed to enter retromer-positive recycling tubules, indicating that the accumulation in endosomes was caused by impaired recycling. APP accumulated in LAMP1-positive endosomes lacking active cathepsins, indicating proteolytic incompetence. Interestingly, the levels of both amyloidogenic and non-amyloidogenic APP-CTFs, but not Aβ, were increased in these compartments. Lastly, we found that these toxic APP-CTFs were secreted by exosomes, small vesicles that form as intraluminal vesicles in the lumen of late endosomes (Bretou et al., 2024). Together, our findings demonstrate that the HOPS complex is required for APP recycling from endosomes to TGN and that depletion of HOPS alters APP processing and secretion, which could contribute to disease propagation.

## Results

### HOPS depletion redistributes APP from the Golgi to endosomes

To investigate the role of the HOPS complex in APP trafficking, we made use of HOPS (VPS18, VPS39 or VPS41) KO HeLa cell lines that were previously made and characterized in our lab (van der Beek et al., 2024). For simplicity, we will generally refer to HOPS KO cells unless we have depleted only one subunit or found a subunit-specific phenotype. HOPS KO cells lack a functional HOPS complex and are impaired in transporting endocytic and autophagic substrates to active lysosomal compartments (Jiang et al., 2014; van der Welle et al., 2021). Moreover, they accumulate enlarged, hybrid early-to-late endosomes, called ‘HOPS bodies’, which indicates a defect in endosomal maturation (van der Beek et al., 2024). To study the effect of HOPS depletion on APP trafficking, we generated an N-terminally mNeonGreen-tagged APP construct (**Fig. 1A**) that we expressed in both WT and HOPS KO HeLa cell lines. Of note, HeLa cells recapitulate characteristics of physiological neuronal APP trafficking and processing in both the secretory and endocytic pathways (Tan and Gleeson, 2019; Januário et al., 2022).

**Figure 1.**
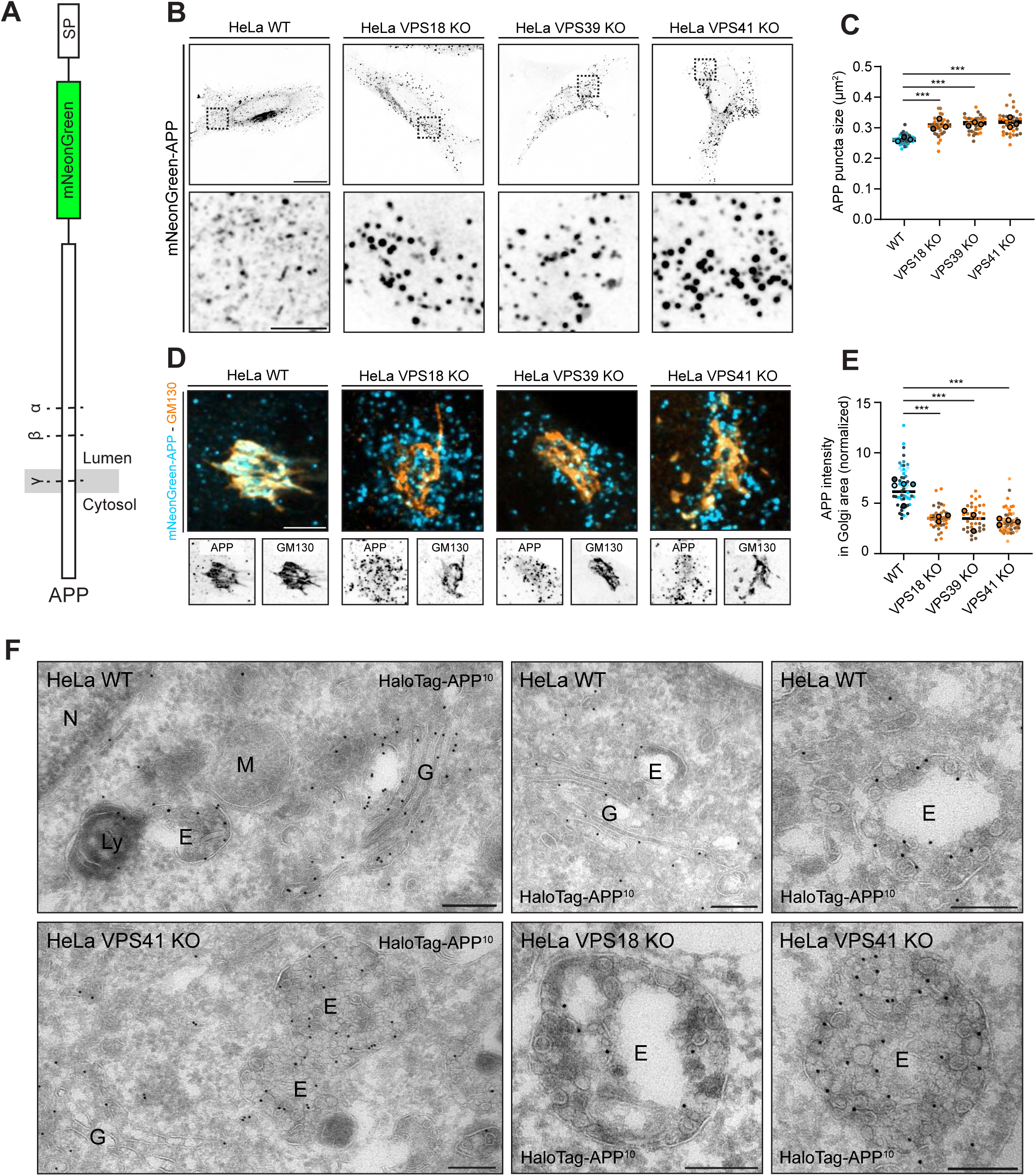
HOPS KO depletes APP from the Golgi area. **A)** Schematic of N-terminal tagged mNeonGreen-APP construct. Cleavage sites by the major secretases are indicated by dashed lines. **B)** Representative images of HeLa WT and HOPS KO cells transiently expressing mNeonGreen-APP. Magnified regions (bottom). **C)** Quantification of average APP puncta size per cell for conditions in (B). **D)** Representative images of HeLa WT and HOPS KO cells expressing mNeonGreen-APP (blue) and stained for GM130 (orange). **E)** Quantification of normalized APP intensity in Golgi region for conditions in (D). **F)** Immuno-EM of HeLa cells expressing HaloTag-APP (15 nm gold particles). N = Nucleus, M = mitochondrion, G = Golgi, Ly = Lysosome, E = Endosome. All scale bars indicate a region of 20 µm (overview), 5 µm (zooms) or 200 nm (EM images). Dots in all dot plots indicate individual cells and lines indicate median value. Mean values of each replicate are indicated in dot blots by bold circles. Ordinary one-way ANOVA was used in (C) followed by Dunnett correction for multiple comparisons. Kruskal-Wallis test was used for (E) followed by Dunn’s correction for multiple comparisons. ns = not significant, *** (P < 0.001).

First, we analyzed the distribution of mNeonGreen-APP in WT HeLa cells by fluorescence microscopy. We found mNeonGreen-APP in the perinuclear area, as well as in small vesicles and tubular compartments dispersed throughout the cytoplasm in WT Hela cells (**Fig. 1B**). This distribution was strikingly changed in all HOPS KO cells. Here, mNeonGreen-APP was largely absent from the perinuclear area and instead accumulated in enlarged vesicles scattered throughout the cytoplasm (**Fig. 1B-C**). The perinuclear pool of mNeonGreen-APP in WT HeLa cells typically reflected Golgi localization. To study this in more detail, we localized mNeonGreen-APP together with GM130, a Golgi marker (**Fig. 1D**). This showed a consistent and drastic depletion of mNeonGreen-APP from the Golgi area in HOPS KO cells (**Fig. 1D-E**).

To study the morphological characteristics of APP-positive compartments, we performed immuno-electron microscopy (immuno-EM) on HeLa WT and HOPS KO cells expressing HaloTag-APP (**Fig. 1F**). HaloTag was used instead of mNeonGreen as it allowed for immuno-EM using an anti-HaloTag antibody. In WT cells, we observed high levels of HaloTag-APP in the TGN, defined as network of membranes at the trans side of the Golgi stack (**Fig. 1F**) (Klumperman, 2011). Additionally, we found HaloTag-APP in tubules and vesicles away from the TGN and in endosomes. By contrast, in HOPS KO cells we found HaloTag-APP mainly in enlarged endosomes with a high intraluminal vesicle (ILV) count (**Fig. 1F**).

Together, these results demonstrate that HOPS KO induces a depletion of mNeonGreen-APP from the Golgi area and redistribution to enlarged endosomes with many ILVs.

### HOPS depletion induces accumulation of APP in late endosomes

To further characterize the nature of the enlarged endosomes accumulating APP in HeLa HOPS KO cells, we performed fluorescence microscopy of APP together with a panel of endo-lysosomal markers. HOPS bodies, observed in HOPS KO cells, are enlarged hybrid compartments positive for both early (EEA1, RAB5, PI3P) and late (RAB7, LAMP1, cathepsin D) endosomal markers (van der Beek et al., 2024), with EEA1 being a distinct marker. Thus, we first examined the occurrence of APP in EEA1-positive endosomes. In HeLa WT cells, around 22% of APP puncta co-distributed with EEA1-positive endosomes. Strikingly, in HOPS KO cells this was reduced to around 12% (**Fig. 2A-B**). Moreover, APP-positive compartments in WT and HOPS KO cells only contained low levels of CI-M6PR, which was previously shown to accumulate in HOPS bodies (**Fig. S1A-B**) (Walia et al., 2024; van der Beek et al., 2024). We then examined colocalization of mNeonGreen-APP with LAMP1, a marker for late endosomes and lysosomes. In WT cells, little colocalization was seen between mNeonGreen-APP and LAMP1 (**Fig. 2C-D**), but in HOPS KO cells this was markedly increased (**Fig. 2C-D**). Furthermore, we found increased colocalization of mNeonGreen-APP with Lysotracker, a marker for acidified compartments (**Fig 2E-F**). However, mNeonGreen-APP was not found in active lysosomes labeled with SiR-Lysosome (SiRLyso), a probe for active cathepsin D (**Fig. 2G-H**).

**Figure 2.**
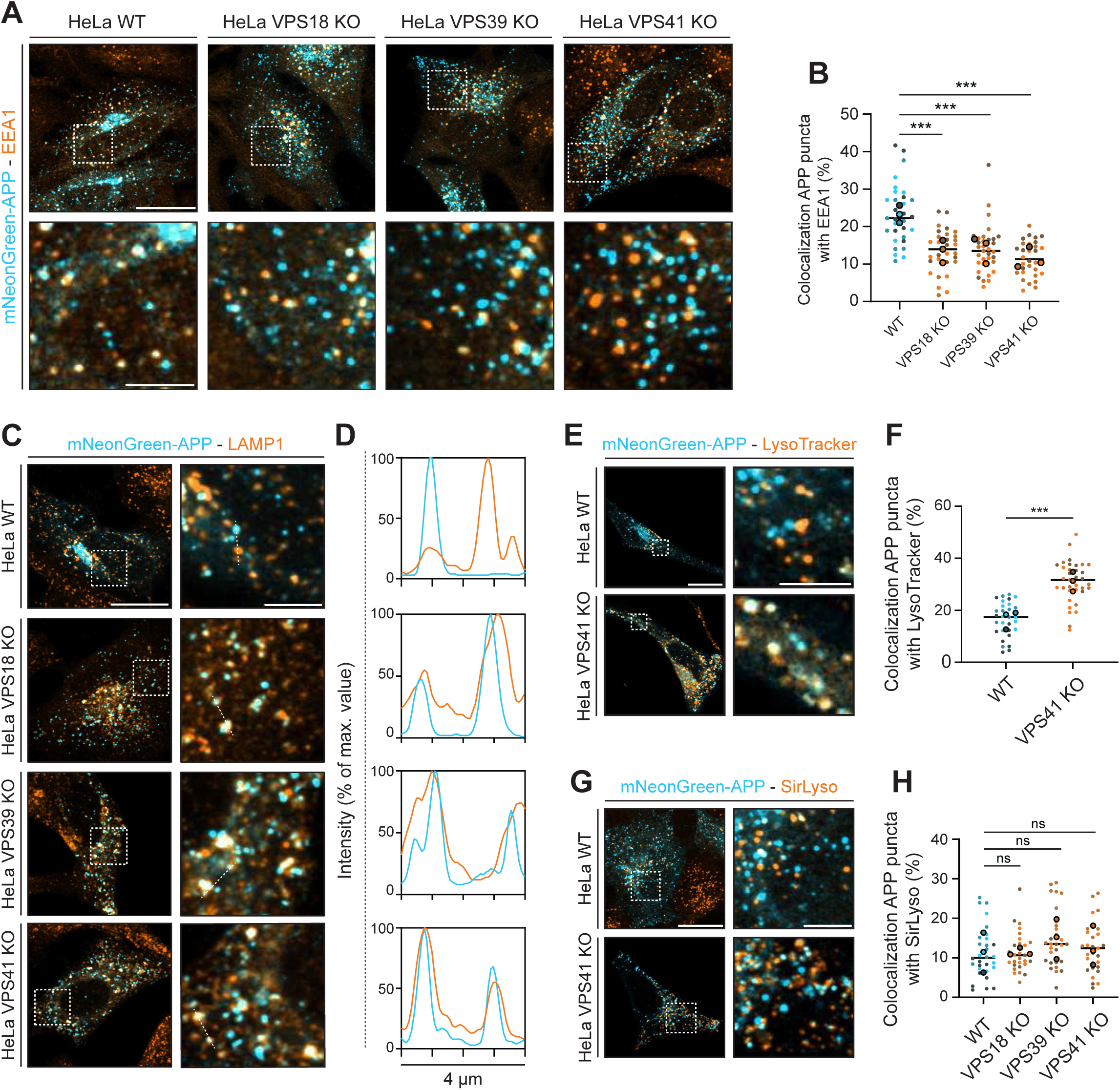
HOPS KO redistributes APP to late endosomes. **A)** Representative images of HeLa WT and HOPS KO cells transiently expressing mNeonGreen-APP (blue) and stained for EEA1 (orange). Magnified regions (bottom). **B)** Percentage of colocalization of APP with EEA1 for conditions in (A). **C)** Representative images of HeLa WT and HOPS KO cells transiently expressing mNeonGreen-APP (blue) and stained for LAMP1 (orange). Magnified regions (right). **D)** Intensity profiles for APP and LAMP1 puncta for conditions in (C). **E)** Representative images of HeLa cells transiently expressing mNeonGreen-APP (blue) and labeled with the probe LysoTracker (orange). Magnified regions (right). **F)** Percentage of colocalization of APP with LysoTracker for conditions in (E). **G)** Representative images of HeLa cells transiently expressing mNeonGreen-APP (blue) and labeled with the probe SirLyso (orange). Magnified regions (right). **H)** Percentage of colocalization of APP with SirLyso for conditions in (G). All scale bars indicate a region of 20 µm (overview) or 5 µm (zooms). Dots in all dot plots indicate individual cells and lines indicate median value. Mean values of each replicate are indicated in dot blots by bold circles. Kruskal-Wallis test followed by Dunn’s correction for multiple comparisons was used in (B) and (H). Unpaired T test was used in (F). ns = not significant, *** (P < 0.001).

We then performed similar colocalization studies with a C-terminal tagged APP construct (APP-mNeonGreen). Like N-terminal tagged mNeonGreen-APP (**Fig. 2E-F**), this construct showed increased colocalization with LysoTracker upon VPS41 KO (**Fig. S1C-D**), indicating that both N- and C-terminal parts of APP accumulated in acidified, late endosome compartments. Furthermore, by immuno-EM we localized C-terminal tagged APP-mNeonGreen (**Fig. S1F**) to morphologically similar compartments as N-terminally tagged APP (**Fig. 1E**), i.e. TGN and endosomes in HeLa WT cells and enlarged endosomes with high ILV content in VPS41 KO cells.

Together these data show that HOPS KO induces redistribution of APP to enlarged, LAMP1-positive compartments with many ILVs, which are acidified but lack catalytic activity, identifying them as late endosomes. Since these endosomes lack EEA1 they are different from the previously described HOPS bodies (van der Beek et al., 2024).

### VPS41 KO impairs endosome-to-TGN recycling of APP

The reduced localization of APP in the Golgi region could be due to defective trafficking through the secretory pathway (Chen et al., 2017) or, alternatively, impaired endosome-to-TGN recycling (Vieira et al., 2010). To address this, we applied the retention using selective hooks (RUSH) tool (Boncompain et al., 2012). In brief, we coupled a streptavidin binding protein (SBP) and a mNeonGreen tag to APP (**Fig 3A**). Co-expression of this with Streptavidin (Strep) coupled to the ER retention signal KDEL, causes the retention of APP at the ER (**Fig. 3A**). Following addition of biotin, APP can be released from the ER, allowing APP trafficking to be visualized in living cells in a synchronized manner (**Fig. 3A**). We expressed the RUSH system for APP in both HeLa WT and HeLa VPS41 KO cells (**Fig 3B**). In both conditions, we found an accumulation of RUSH-APP in the Golgi area approximately at 20 minutes after ER release, with a peak intensity around 40 minutes and a drop in intensity after 60 minutes release (**Fig. 3C**). Notably, we found no significant difference in the kinetics of RUSH-APP release from the Golgi region between WT and KO cells (**Fig 3D; Video S1),** indicating that VPS41 does not alter APP sorting from the TGN. We then assessed the effect of HOPS depletion on endosomal recycling of APP by monitoring RUSH-APP at 1 and 8 hours after release from the ER (**Fig. 3E**). A defect in recycling would cause a further drop in intensity in the Golgi region after 8 hours of biotin addition. In HeLa WT cells, we found a modest reduction in RUSH-APP intensity in the Golgi area after 8 hours of release, but overall the signal remained abundantly present (**Fig. 3E-F**). This is consistent with previous studies showing continuous endosome-to-TGN recycling of APP (Vieira et al., 2010). By contrast, VPS41 KO cells showed a marked reduction of the RUSH-APP signal in the Golgi area after 8 hours, indicating a defect in recycling from endosomes to the TGN (**Fig. 3E-F**). This observation is consistent with the immuno-fluorescence data obtained from HOPS KO cells expressing mNeonGreen-APP (**Figure 1D, E**).

**Figure 3.**
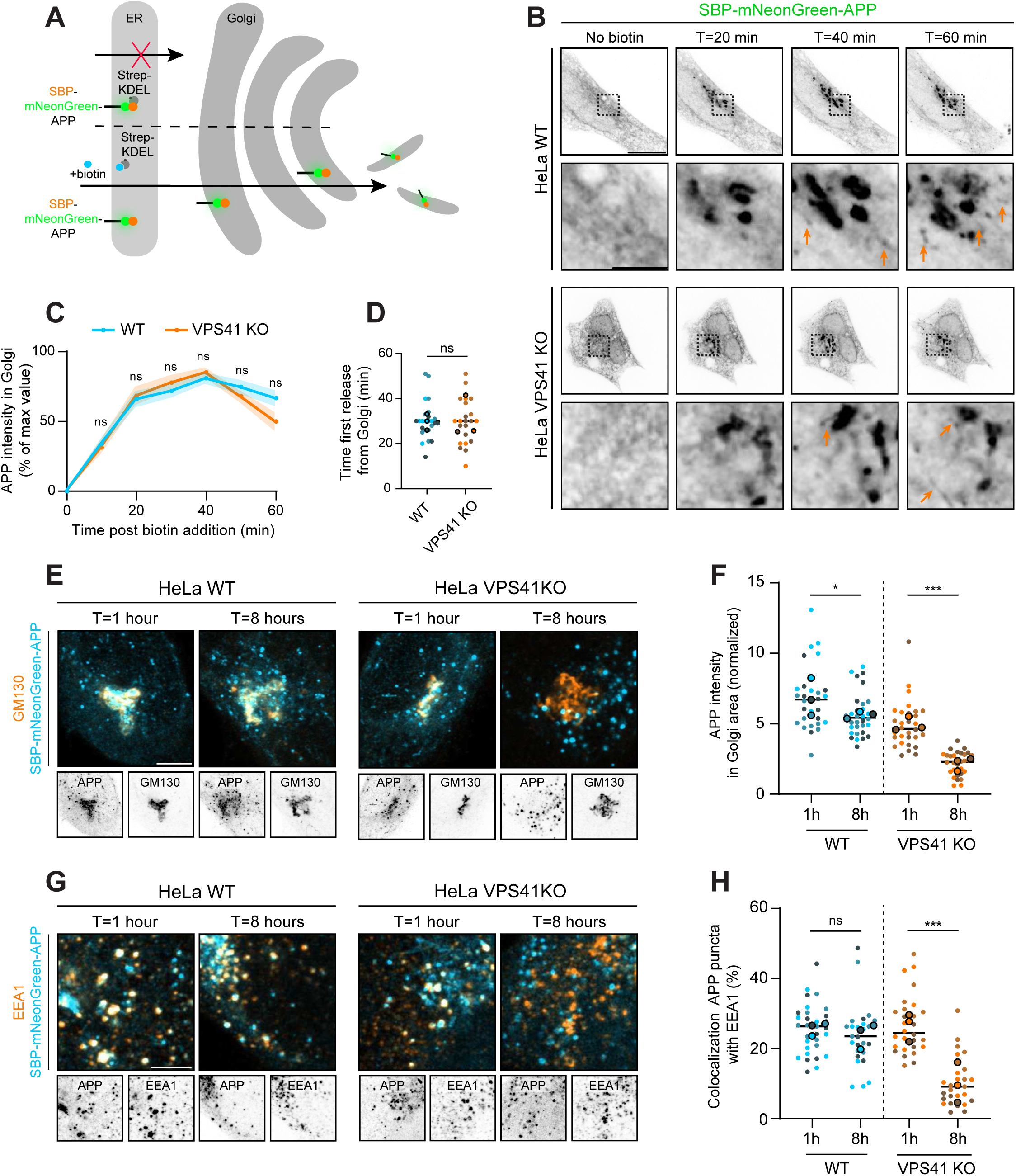
VPS41 depletion impairs APP endosome-to-TGN recycling. **A)** Schematic of the RUSH system for retention and release of APP from the ER. **B)** Representative still images of videos from HeLa WT and VPS41 KO cells expressing SBP-mNeonGreen-APP. Cells were imaged every minute for 60 min. Magnified regions (bottom), orange arrows point to release events from the Golgi. **C-D)** Quantification of APP intensity in the Golgi area over time **(C)**, and time of the first detected APP tubule being released from the Golgi **(D)**, from conditions in (B). **E)** Representative images of HeLa WT and VPS41 KO cells expressing SBP-mNeonGreen-APP (blue) and stained for GM130 (orange). **F)** Quantification of normalized APP intensity in Golgi area upon 1h and 8h ER-release, from conditions in (E). **G)** Representative images of HeLa WT and VPS41 KO cells expressing SBP-mNeonGreen-APP (blue) and stained for EEA1 (orange). **H)** Percentage of colocalization of APP with EEA1 upon 1h and 8h ER-release, from conditions in (G). Time points (T=) indicate time after biotin addition in (B), (E), and (G). Scale bars indicate a region of 20 µm in overview and 5 µm in zooms. Dots in all dot plots indicate individual cells and lines indicate median value. Mean values of each replicate are indicated in dot blots by bold circles. Data is visualized as mean value plus SEM in (C). Ordinary two-way ANOVA followed by Tukey’s correction for multiple comparisons was used in (C), (F) and (H). Mann-Whitney test was used in (D). ns = not significant, * (P = 0.033), *** (P < 0.001).

Early endosomes – also called sorting endosomes – are important sites for the formation of tubules that recycle cargo to the TGN or plasma membrane (Burd and Cullen, 2014). For this reason, we examined RUSH-APP colocalization with the early endosome marker EEA1 at different time points (**Fig. 3G**). After 1 hour release, comparable levels of RUSH-APP were detected in EEA1-labelled endosomes of both WT and VPS41 KO cells (**Fig. 3G-H**). Interestingly, while this percentage remained stable in WT cells after 8 hours of release, it was markedly reduced in VPS41 KO cells at this later time point (**Fig. 3G-H**). These results are consistent with our finding that in HOPS KO cells APP accumulates in late endosomes rather than in EEA1-positive HOPS bodies (**Fig. 2**). Additionally, after 8 hours release, the percentage of cells that contained RUSH-APP-positive tubules dropped drastically in HeLa VPS41 KO but not WT cells, suggesting loss of APP from recycling tubules rather than from biosynthetic, TGN-derived tubules (**Fig. S2A-B**).

Together these data indicate that in the absence of VPS41, APP trafficking from TGN to EEA1-positive endosomes is normal. However, APP recycling from endosomes to the TGN is impaired, resulting in accumulation of APP in late endosomes.

### HOPS depletion prevents recruitment of APP into VPS35 recycling tubules

Since the RUSH experiment indicated a defect in APP recycling from endosomes, we used the resolution of immuno-EM to assess colocalization of APP with VPS35, a component of the retromer complex known to be involved in the recycling of APP (**Fig. 4A**) (Vieira et al., 2010). Previous EM studies have shown that VPS35 is typically associated with recycling tubules (Butkovič et al., 2024). In WT HeLa cells we could readily identify these VPS35-positive recycling tubules and by double-labeling for HaloTag-APP show that they contained APP (**Fig. 4A**). Interestingly, in VPS41 KO cells, VPS35-positive recycling tubules were still formed from APP-positive endosomes, but they now generally lacked APP (**Fig. 4A**). To validate and extend these data we used the mNeonGreen-APP construct to assess colocalization with VPS35 and the VPS35-interactor SNX1 by fluorescence microscopy (Zhang et al., 2018). In WT cells, we observed around 37% and 42% colocalization of mNeonGreen-APP with VPS35 and SNX1, respectively (**Fig. 4B-E**) This level of colocalization was drastically reduced (around 16% for VPS35 and 18% for SNX1) in HOPS KO cells (**Fig. 4B-E**). To exclude that this reduction was due to decreased protein levels of VPS35, we performed immunoblotting on whole cell lysates. This showed that VPS35 levels were not altered in HOPS KO compared to WT cells (**Fig. 4F**).

**Figure 4.**
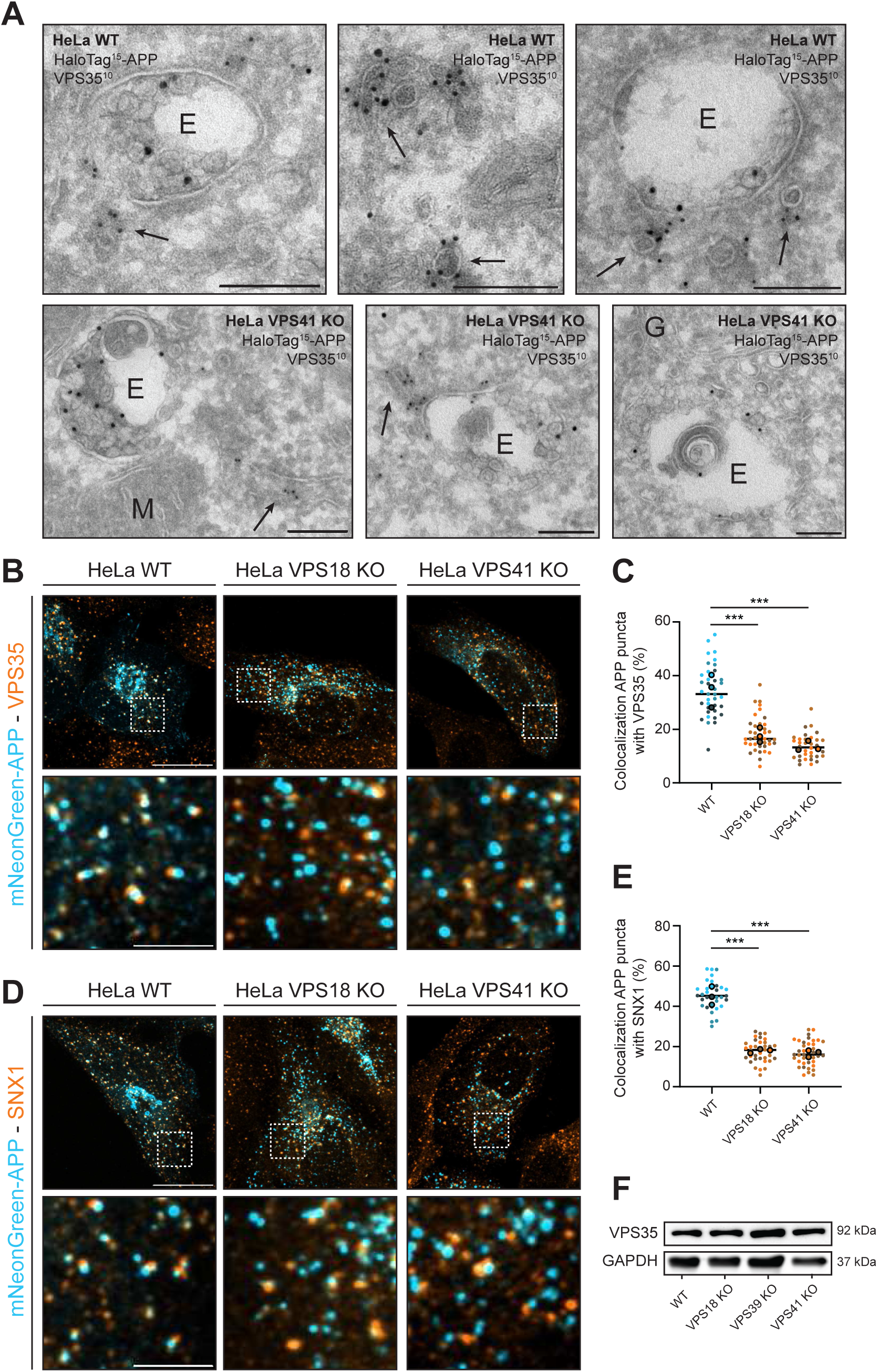
HOPS KO impairs APP entry into retromer-positive recycling tubules. **A)** Immuno-EM of HeLa WT (top) and VPS41 KO (bottom) cells expressing HaloTag-APP (15 nm gold) and immuno-labelled for retromer subunit VPS35 (10nm gold). Arrows indicate recycling tubules. E= endosome, M = mitochondrion, G = Golgi. **B, D)** Representative images of HeLa WT and HOPS KO cells transiently expressing mNeonGreen-APP (blue) and stained for VPS35 (orange) **(B)** or SNX1 (orange) **(D)**. Respective magnified regions (bottom). **C, E)** Percentage of APP colocalization with VPS35 **(C)** or SNX1 **(E)**, from conditions in (B) and (D). **F)** Western blot for endogenous VPS35 and GAPDH in HeLa WT and HOPS KO cells. Scale bars indicate 20 µm in overview, 5 µm in zooms, and 200 nm in EM images. Dots in all dot plots indicate individual cells and lines indicate median value. Mean values of each replicate are indicated in dot blots by bold circles. Kruskal-Wallis test followed by Dunn’s correction for multiple comparisons was used in (C). Ordinary one-way ANOVA followed by Dunnett correction for multiple comparisons was used in (E). ns = not significant, *** (P < 0.001).

Together, these results show that APP fails to enter VPS35- and SNX1-positive recycling tubules that form from APP-positive endosomes in HOPS KO cells. This is consistent with the RUSH-APP experiments showing that APP does not recycle from endosomes to TGN in VPS41 KO cells (**Fig. 3E-F**). Because of this failed recycling, APP accumulates in LAMP1-positive late endosomes **(Fig. 2**).

### HOPS depletion increases APP and BACE1 colocalization and induces APP-CTF accumulation

Endosomes are major sites for amyloidogenic APP processing by the β-secretase BACE1 (see for a schematic of APP processing, **Fig 5A**) (Das et al., 2015; Park et al., 2022). Since HOPS KO induced redistribution of APP from TGN to late endosomes, we wondered whether these might contain BACE1 as well. In physiological circumstances BACE1 is mainly present in recycling endosomes (Das et al., 2013; Tan et al., 2020) and in HeLa WT cells, we found that around 20% of mNeonGreen-APP colocalized with BACE1-mScarlet (**Fig. 5B, C**). Strikingly, this percentage increased to around 40% in HOPS KO cells (**Fig. 5B, C**), indicating that upon HOPS depletion APP and BACE1 redistribute to the same endosomal compartments, resulting in enhanced colocalization.

**Figure 5.**
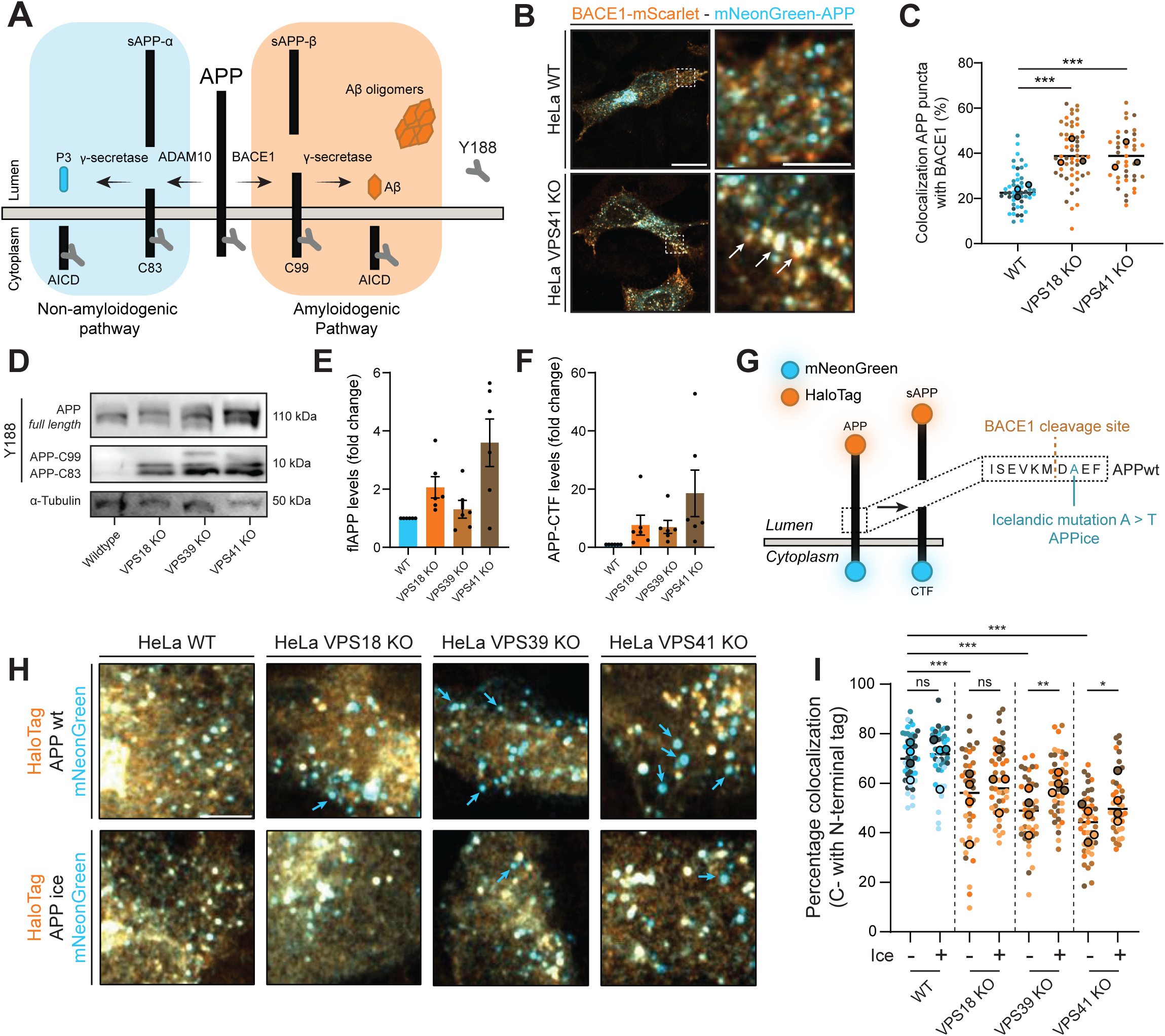
HOPS depletion increases APP-CTF (C83 and C99) levels. **A)** Schematic of both non-amyloidogenic and amyloidogenic processing of APP and its derived fragments. **B)** Representative images of HeLa WT and VPS41 KO cells transiently expressing the amyloidogenic secretase BACE1-mScarlet and mNeonGreen-APP. Arrows indicate APP and BACE1 in the same compartments. Magnified region (right). **C)** Quantification of the colocalization between APP and BACE1 as in (B). **D)** Western blot of endogenous full-length APP (flAPP) and APP-CTFs using the C-terminal antibody Y188 (binding site indicated in A). **E-F)** Fold change of normalized flAPP **(E)**, and normalized APP-CTF **(F)** levels in HOPS KO cells compared to WT cells. Dots indicate individual replicates. APP-CTF levels were normalized to α-tubulin levels. **G)** Schematic of double-tagged HaloTag-APP-mNG. Increased processing of APP would result in decreased colocalization of C- and N-terminal tag. Amino acid sequence of APP_wt_ and APPi_ce_ (A673T) are indicated on the right. Icelandic mutation is protective for BACE1 cleavage. **H)** Representative images of HeLa WT and HOPS KO cells expressing either double-tagged APP_wt_ or APP_ice_ constructs. Blue arrows indicate compartments only containing C-terminally tagged APP (APP-CTF). **I)** Quantification of the colocalization between C- and N-terminal APP. All scale bars indicate a region of 20 µm (overview) or 5 µm (zooms). Dots in all dot plots indicate individual cells and lines indicate median value. Mean values of each replicate are indicated in dot blots by bold circles. Data is visualized as mean + SEM in (E) and (F). Ordinary one-way ANOVA followed by Dunnett correction for multiple comparisons was used in (C). Ordinary two-way ANOVA followed by Tukey’s correction for multiple comparisons was used in (I). ns = not significant, * (P = 0.033), ** (P = 0.002), *** (P < 0.001).

BACE1 mediates APP processing most efficiently at low pH (Vassar et al., 1999). Since we showed in VPS41 KO cells that APP relocates to LysoTracker-positive (acidified) late endosomes, where APP shows enhanced colocalization with BACE1, we wondered if these could be sites for amyloidogenic processing. To address this, we performed immunoblotting on whole cell lysates. Using antibody Y188, which targets the C-terminal tail of APP (indicated in schematic in **Fig. 5A**) (Guo et al., 2011), we could detect full-length APP (flAPP) and its derivatives (the APP-CTFs C83, C99 and AICD) and study APP processing in WT and HOPS KO cells. Since the cytoplasmic AICD fragment is rapidly cleared by cytoplasmic metalloproteases rather than by lysosomal degradation, we focused on C83 and C99 for further experiments (Edbauer et al., 2002). In WT cells, we could readily detect flAPP but no C83 or C99, suggesting that these CTFs upon formation are efficiently degraded in lysosomes or further processed by the γ-secretase complex (**Fig. 5D-F**). Interestingly, in HOPS KO cells we observed a slight increase in flAPP levels and a striking increase in APP-CTFs, including the amyloidogenic C99 fragment (**Fig. 5D-F**). Thus, HOPS depletion results in the accumulation of APP-CTFs.

To further comprehend the effect of HOPS depletion on APP processing, we used a previously reported dual-tagged APP construct, with HaloTag on the N-terminus and mNeonGreen at the C-terminus (**Fig. 5G**) (Sannerud et al., 2011; Villegas et al., 2013; Hitschler and Lang, 2022; Januário et al., 2022). A decrease in colocalization between the C- and N-terminal tags indicates increased processing. Consistent with our immunoblot results, we found a high level of colocalization in HeLa WT cells (**Fig. 5H-I**), whereas HOPS KO cells displayed numerous fluorescent puncta positive for the C-terminal tag but lacking N-terminal signal. We speculated that the absence of N-terminal APP may result from release of sAPP into the extracellular space, either via APP cleavage at the plasma membrane or through endosomal secretion (**Fig. 5A**) (Hitschler and Lang, 2022). Collectively, these findings indicate increased processing of APP in HOPS KO cells, which is consistent with the substantial accumulation of APP-CTFs observed by immunoblotting (**Fig. 5D**).

To distinguish between amyloidogenic and non-amyloidogenic processing we next used a dual-tagged construct of the APP Icelandic mutation (A673T), henceforth called APP_ice_ (**Fig. 5G**), which is known to reduce amyloidogenic processing (Maloney et al., 2014; Das et al., 2015). By comparing APP_wt_ to APP_ice_ processing we assessed the relative contribution of amyloidogenic versus non-amyloidogenic processing in WT versus HOPS KO cells. In HeLa WT cells, the level of colocalization between both tags was similarly high for APP_wt_ and APP_ice_, indicating little amyloidogenic processing (**Fig. 5H, I**). By contrast, expression of APP_ice_ in VPS39 or VPS41 KO cells showed a significant increase in colocalization between the C- and N-terminal tags (**Fig. 5H, I**), although this remained lower than in WT cells. These data suggest that HeLa WT cells perform little amyloidogenic processing, while HOPS KO cells accumulate both amyloidogenic and non-amyloidogenic APP-CTFs.

We conclude from these experiments that HOPS depletion results in increased APP-CTF levels, which is partially due to amyloidogenic processing by BACE1. Besides the striking increase in APP-CTFs, we also found increased levels of flAPP, which is probably due to decreased degradation in lysosomes, a known phenotype in cells lacking the HOPS complex.

### HOPS disruption induces endosomal accumulation of APP in the somatodendritic domain of rat hippocampal neurons

Since neurons are especially susceptible to lysosomal dysfunction and given that formation of amyloid plaques occurs in the brain (Chou et al., 2025), we next wanted to assess the role of HOPS in APP trafficking in rat hippocampal neurons. Neurons are highly polarized cells divided in somatodendritic and axonal domains (Britt et al., 2016; Bentley and Banker, 2016). AD is characterized by specific defects in both subdomains, including endo-lysosomal traffic jams, severe loss of dendritic spines and axon degeneration (Gowrishankar et al., 2015; Small et al., 2017; Subramanian et al., 2020; Salvadores et al., 2020). We used rat hippocampal neurons at day-in-vitro 8 (DIV8), which are fully polarized and allowed studying the effect of HOPS depletion in the respective subdomains.

To deplete VPS41 in neurons, we expressed 4 different shRNAs targeting VPS41, for 4 days. Immuno-fluorescence labeling of endogenous VPS41 at DIV8 showed that the staining significantly decreased in these neurons as compared to control conditions (**Fig. S3A-B**). However, knockdown of VPS41 in these primary neurons also induced massive neurite fragmentation and cell death (**Fig. S3C-D**). Although these results show the importance of the HOPS complex in neuronal maintenance, this prevented us from studying its contribution to APP trafficking.

As an alternative approach to disrupt HOPS complex formation in neurons, we transfected cells with ORF3a-mCherry. ORF3a, a SARS-CoV2 accessory protein, binds to the VPS39 subunit of the HOPS complex, thereby impairing HOPS functioning and resulting in impaired lysosomal fusion events (schematic in **Fig. 6A**) (Miao et al., 2020; Walia et al., 2024). First, we validated the effect of ORF3a-mCherry expression on mNeonGreen-APP distribution in HeLa cells (**Fig. 6B, C**). After 24h, ORF3a expression led to a depletion of APP from the Golgi area, similar as seen in HeLa HOPS KO cells, (**Fig. 6B, C**). Subsequently, we expressed ORF3a-mCherry or mCherry control (empty mCherry) for 24h in neurons, after which we labeled for endogenous APP with the Y188 antibody. In control neurons, we found a significant overlap between endogenous APP and the Golgi marker GM130 (**Fig. 6D, E**). In neurons expressing ORF3a-mCherry, however, APP was depleted from the GM130-positive Golgi area and accumulated in vesicular compartments (**Fig. 6D, E**). To ensure that the effect of ORF3A on APP distribution was indeed HOPS-dependent, we next introduced two mutations in the ORF3a-mCherry construct (S171E or W193R), which impair binding to VPS39 (D. Chenet al., 2021; Walia et al., 2024). Expression of these mutants failed to deplete APP from the Golgi region (**Fig. 6D, E**), indicating that the effect of ORF3a on APP distribution in neurons is HOPS dependent.

**Figure 6.**
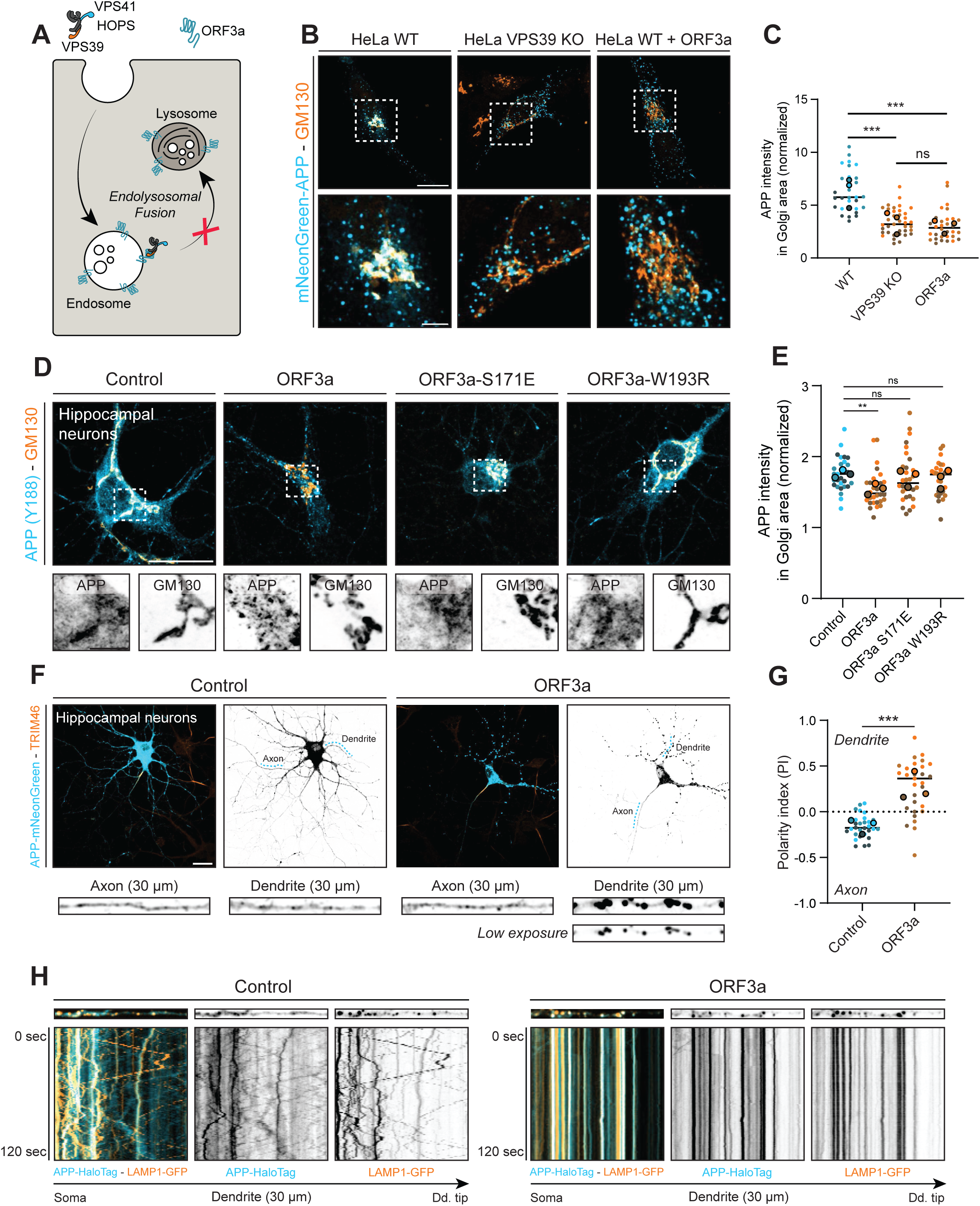
HOPS disruption induces accumulation of APP in stationary LAMP1-positive endosomes in the somatodendritic domain of rat primary neurons. **A)** Simplified mode of action of SARS-CoV2 accessory protein ORF3a. ORF3a is a transmembrane protein that can directly bind to VPS39, leading to the inhibition of HOPS functioning. **B)** Representative images of mNeonGreen-APP (blue) localization in HeLa WT, VPS39 KO and ORF3a expressing cells. Golgi region was stained using the Cis-Golgi marker GM130. Magnified region (bottom). **C)** Quantification of APP intensity in Golgi area from conditions in (B). Data for HeLa WT and VPS39 KO cells were re-used in Fig. 1E. **D)** Representative images of DIV8 hippocampal neurons transfected on DIV7 with either mCherry (control), ORF3a-mCherry, ORF3a(S171E)-mCherry or ORF3a(W193R)-mCherry. Neurons were stained for endogenous APP (Y188) (blue) and GM130 (orange). Magnified region (bottom). **E)** Quantification of normalized APP intensity in Golgi region in Control and ORF3a expressing neurons as in (D). **F)** Representative images of DIV8 neurons co-transfected on DIV7 with C-terminally tagged APP-mNeonGreen and control (mCherry) or ORF3-mCherry, and labeled for the axon initial marker TRIM46. Zoom of axon and dendrite segments (bottom) **G)** Quantification of polarity index in control and ORF3a transfected neurons. **H)** Stills (top) and kymographs (bottom) of live neurons expressing APP-HaloTag (blue) and LAMP1-GFP (orange) in control and ORF3a expressing neurons. Dd. tip = dendrite tip. All scale bars indicate a region of 20 µm (overview) or 5 µm (zooms). Dots in all dot plots indicate individual cells and lines indicate median value. Mean values of each replicate are indicated in dot blots by bold circles. Kruskal-Wallis test followed by Dunn correction for multiple comparisons was used in (C) and (E). Welch’s t test was used in (G). ns = not significant, ** (P = 0.002), *** (P < 0.001).

Transport of APP to dendrites and the axon, with a preferential sorting towards the axon, plays a key role in axon and dendrite growth, synapse formation and maintenance (Hoe et al., 2010; van der Kant and Goldstein, 2015). To assess a potential effect of HOPS disruption on the distribution of APP over distinct neuronal subdomains, we expressed C-terminally tagged APP-mNeonGreen together with ORF3a-mCherry or mCherry control (**Fig. 6F, G**). In this case we used overexpressed APP to facilitate identification of axons and dendrites originating from the same cell, thereby minimizing false-positive signals caused by overlapping neurites from neighboring cells. We calculated the polarity index of APP distribution (PI = [intensity dendrite − intensity axon] / [intensity dendrite + intensity axon], in which PI = 0, unpolarized; PI > 0, dendritic; PI < 0, axonal) for each condition (Kapitein et al., 2010a). In control cells, APP was slightly polarized towards the axonal domain (**Fig. 6F-G**), as previously reported (Allinquant et al., 1994). To our surprise, however, in ORF3a expressing cells, APP accumulated in the somatodendritic domain, leading to a drastic shift in the polarity index towards dendrites (**Fig. 6F-G**). Of note, we could still observe APP-positive compartments in the axon, indicating that the shift in polarity index was caused by an accumulation of APP and likely APP-CTFs in dendrites rather than a reduction in the axonal domain.

To investigate the dynamics and identity of dendritic APP-positive compartments in control versus HOPS impaired neurons, we co-expressed APP-HaloTag together with LAMP1-GFP and studied their transport kinetics by live cell imaging. In control neurons, APP and LAMP1 mostly localized to separate compartments, moving in an antero- and retrograde fashion along the dendrites (**Fig. 6H; Video S2**). Strikingly, ORF3a expression led to a drastic increase in colocalization of APP and LAMP1 in compartments that, surprisingly, were mostly stationary (**Fig. 6H; Video S2**). To determine whether this change in localization and dynamics was specific for APP-CTFs, flAPP, or both, we conducted further analysis with the N-terminal tagged HaloTag-APP construct (**Fig. S3E**), which labels flAPP as well as sAPP. Similar to C-terminal tagged APP (**Fig. 6H**), HaloTag-APP in control cells localized to mobile, LAMP1-negative compartments and in ORF3a expressing cells to stationary LAMP1-positive compartments (**Fig. S3E**; **Video S3**). The data are consistent with our findings in HeLa cells showing that HOPS KO causes accumulation of both APP C- and N-termini in LAMP1-positive late endosomes (**Fig. 2C-D; Fig. S1C-E)**. The local accumulation of APP-CTFs in static dendritic endo-lysosomes could possibly contribute to dendritic spine loss, which is an early hallmark in AD patients (Hoe et al., 2010; Luo et al., 2025).

In conclusion, we find that HOPS disruption by ORF3a in primary neurons causes a depletion of endogenous APP from the Golgi area, and leads to a redistribution and accumulation of APP and likely CTFs in static, LAMP1-positive endosomes that are predominantly present in the somatodendritic domain.

### HOPS disruption in neurons increases APP-CTF levels but not Aβ42 production

To study the effect of HOPS inhibition on APP processing in neurons, we optimized lentiviral transduction of ORF3a in rat cortical neurons at DIV14-15. Neurons were transduced on DIV7 with ORF3a-mCherry or control mCherry, reaching a transduction efficiency of 80-90% (**Fig. S4A**). Cells were harvested and whole cell lysates were prepared for immunoblotting with antibody Y188 (recognizing flAPP and APP-CTFs). Similar as in HeLa HOPS KO cells, we found that HOPS disruption in neurons increased APP-CTF levels (**Fig. 5D-F**; **Fig. 7A-C**). As a positive control we treated neurons with the ɣ-secretase inhibitor DAPT, which prevents further processing of APP-CTFs (Tan et al., 2020; Burrinha et al., 2021). Of note, BACE1 and γ-secretase subunit PSEN2 levels remained unaffected by ORF3a expression, indicating that the increase in APP-CTFs levels was not caused by increased BACE1 or decreased γ-secretase expression (**Fig. S4B-E**). Additionally, we transduced neurons with the double-tagged APP construct (HaloTag-APP-mNG) in order to visualize APP processing (**Fig. 7D-E**). Consistent with the immunoblotting data (**Fig. 7A-C**) and with our findings in HeLa cells (**Fig. 5I-H**), we found reduced colocalization of C- and N-terminal tag in ORF3a expressing neurons, indicating increased processing (**Fig. 7D-E**).

**Figure 7.**
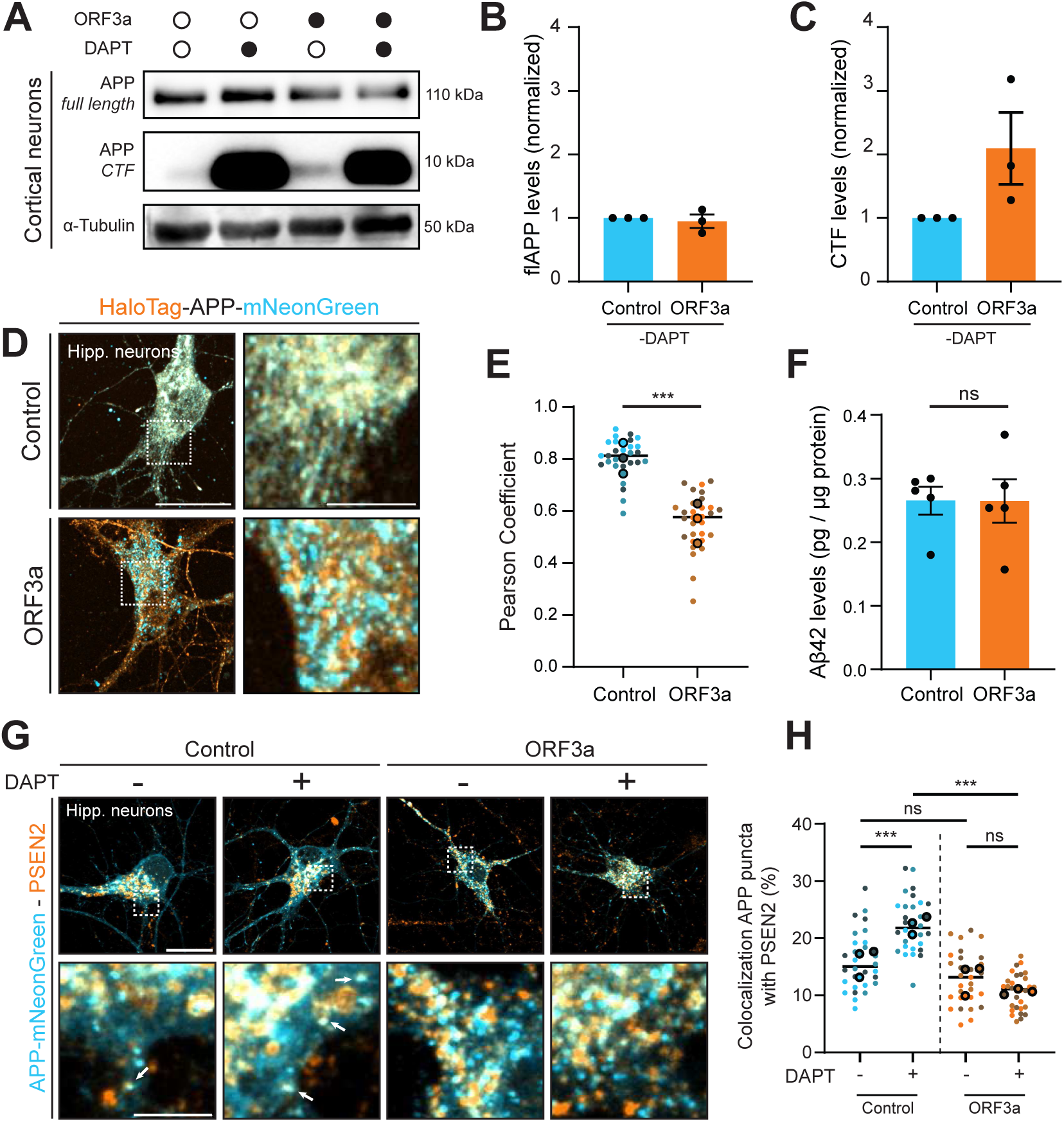
HOPS impairment in primary neurons increases APP-CTF levels. **A-C)** Cortical neurons transduced on DIV7 with lentivirus containing mCherry or ORF3a-mCherry were treated for 16h with 25 µM γ-secretase inhibitor or DMSO control on DIV14 prior to lysis and western blot for APP (Y188) on DIV15. Fold change compared to mCherry (control) in protein levels of flAPP **(B)** and APP-CTFs **(C)** from DMSO-treated cells. Individual dots indicate replicates. **D)** Representative images of DIV8 hippocampal neurons transduced on DIV4 with lentivirus containing HaloTag (orange)-APPwt-mNeonGreen (blue) plus either mCherry (control) or ORF3a-mCherry (mCherry label not shown). Magnified regions (right). **E)** Pearson correlation of APP C- and N-terminal tag in control and ORF3a treated neurons. **F)** ELISA measurement of Aβ42 of cell lysates as indicated in (A). **G)** Representative images of DIV8 hippocampal neurons transiently expressing C-terminally tagged APP-mNeonGreen (blue) with either mCherry (control) or ORF3a-mCherry, and stained for the γ-secretase subunit PSEN2 (orange). Neurons were treated for 16h with 25 µM γ-secretase inhibitor to block γ-secretase activity or DMSO control, prior fixation. **H)** Quantification of APP-mNeonGreen colocalization with endogenous PSEN2. All scale bars indicate a region of 20 µm (overview) or 5 µm (zooms). Dots in all dot plots indicate individual cells and lines indicate median value. Mean values of each replicate are indicated in dot blots by bold circles. Data are presented as mean values ± SEM in (B), (C), and (F). Mann-Whitney test was used in (E). Welch’s t test was used in (F). Ordinary two-way ANOVA followed by Tukey’s correction for multiple comparisons was used in (H). ns = not significant, * (P > 0.033), *** (P < 0.001).

To assess whether accumulation of APP-CTF fragments in ORF3A transfected neurons led to increased Aβ42 production, we used an ELISA-based approach to measure Aβ42 levels (Gowrishankar et al., 2017). To test the specificity of the kit, we measured Aβ42 levels in neurons treated with and without DAPT (**Fig S4F-G**). DAPT treatment resulted in decreased Aβ42 levels, ensuring the validity of this test. To our surprise, however, we did not see an increase in Aβ42 levels in ORF3A transfected neurons (**Fig. 7F**), indicating no further processing of APP-CTFs by the γ-secretase complex.

PSEN2, the catalytic subunit of the γ-secretase complex, is located in late endosomes and lysosomes (Sannerud et al., 2016). Since HOPS is required for endosome-lysosome fusion, we reasoned that the HOPS complex could be necessary for targeting APP-CTFs to PSEN2-positive compartments. To test this, we expressed C-terminally tagged APP-mNeonGreen and investigated colocalization with PSEN2 upon ORF3a expression (**Fig. 7G**). In both control and ORF3a expressing neurons, we found around 14% colocalization between APP and PSEN2 (**Fig. 7G-H**). We then additionally treated neurons with DAPT to enhance detection of APP-CTFs. In control cells, DAPT treatment caused an increase in colocalization of APP-mNeonGreen with PSEN2, indicating efficient targeting of APP-CTFs to PSEN2-containing endo-lysosomes (**Fig. 7G-H**). By contrast, DAPT treatment in ORF3a treated cells did not increase colocalization of APP-CTFs with PSEN2 (**Fig 7G-H**), indicating that HOPS is required for APP-CTF targeting to PSEN2-positive compartments.

Together, our data indicated that HOPS disruption promotes accumulation of toxic APP-CTFs in neurons, but not the production of Aβ42, since the delivery of APP-CTFs to PSEN2-positive compartments is HOPS dependent.

### HOPS disruption increases secretion of APP-CTFs by exosomes

Recently it was shown that inhibition of fusion between endosomes and lysosomes through VPS39 depletion, promotes fusion of late endosomes with the plasma membrane, resulting in the secretion of extracellular vesicles (EVs) (Shelke et al., 2023). Since we found that HOPS disruption induces accumulation of flAPP and APP-CTFs in late endosomes, we asked if flAPP and APP-CTFs could be secreted via EVs. To obtain sufficient numbers of EVs, we isolated EVs from HeLa WT and HOPS KO culture medium by ultracentrifugation and size exclusion chromatography (SEC) (schematic in **Fig. S5A**). To validate our EV isolation procedure, we immunoblotted for calnexin (ER marker) and CD63 (a marker for late endosome-derived EVs) in all fractions isolated from HeLa VPS41 KO cells. Calnexin was only detected in cell lysates, while CD63 was found in both lysates and EVs, confirming the purity of our EV isolation procedure (**Fig. S5B)**.

Experiments were performed in HeLa WT cells and KO for VPS18, VPS39 and VPS41. We first assessed by quantitative immunoblotting the abundance of CD63 in all samples. Strikingly, all HOPS KO lines showed elevated CD63 levels in both cell lysate and EV fractions, with the most prominent increase in the EV fraction of VPS41 KO cells (**Fig. 8A, C**). Subsequentially, we specifically determined the flAPP and APP-CTF levels per condition. This showed that in HOPS KO cell lysates, both flAPP and APP-CTFs were increased compared to WT cells, with a most drastic increase in VPS41 KO cells (**Fig. 8B**). Surprisingly, APP-CTFs were present in all HOPS KO cells and were particularly enriched in EVs from VPS41 KO cells (**Fig. 8B, D**). Notably, flAPP was absent from all EV fractions, including those from VPS41 KO cells (**Fig. 8B**).

**Figure 8.**
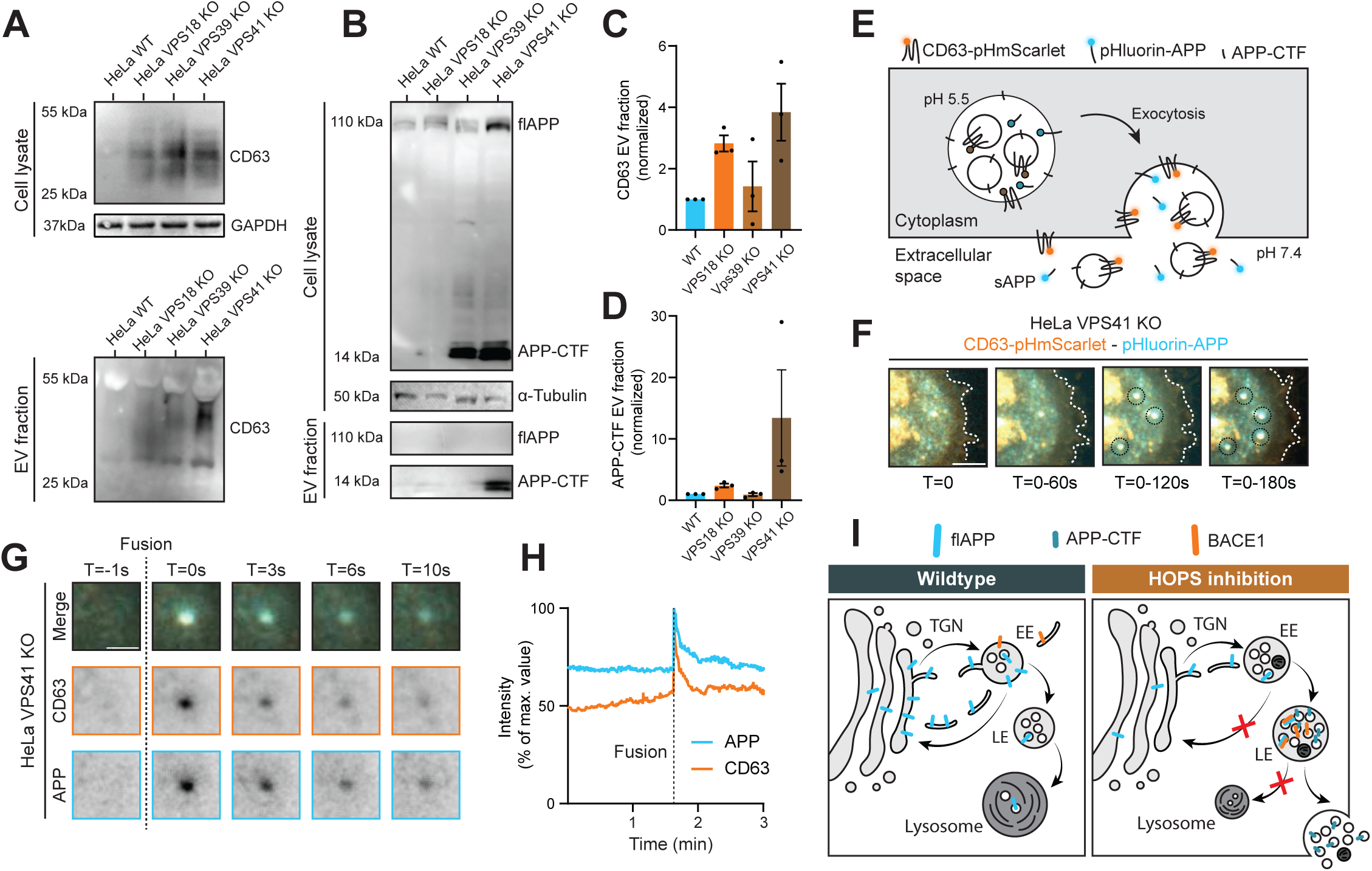
HOPS KO increases secretion of APP-CTFs by extracellular vesicles. **A)** Western blot analysis of CD63 in cell lysates and EV fraction in HeLa WT and HOPS KO cell lines. GAPDH was used as loading control **B)** Western blot analysis of flAPP and APP-CTFs of HeLa WT and HOPS KO cells using C-terminal directed APP antibody (Y188). α-tubulin was used as loading control. **C-D)** Quantification of CD63 and APP-CTF levels in EV fraction. APP-CTF levels are normalized to loading control levels in cell lysates. Dots indicate individual replicates. **E)** Schematic of combined visualization of APP together with exosomal marker CD63. Both proteins are tagged with pH sensitive reporters, which is quenched in the endolysosomal environment. Fusion with the PM would result in an increase in fluorescent signal. **F)** TIRF microscopy of pHluorin-APP (blue) and CD63-pHmScarlet (orange) in VPS41 KO cells. Images were acquired every 0.33 seconds for 3 minutes. Maximum projection indicates the dual positive fusion events (indicated by dashed circles). Scale bar indicates 5 µm. **G)** Dynamics of single fusion event in similar video as in (F). Time is indicated as seconds respective to fusion event. **H)** Intensity dynamics profiles of fusion event in (G). Time of fusion is indicated by the dotted line in the graph. **I)** Proposed model of APP trafficking and processing alterations upon HOPS disruption. EE = early endosome / HOPS body, LE = late endosome.

Next, to visualize EV secretion in live cells, we performed total internal reflection fluorescence (TIRF) microscopy of VPS41 KO cells transfected with CD63 and APP, both fused to pH-sensitive probes (pHmScarlet and pHluorin, respectively) (schematic in **Fig. 8E**) (Verweij et al., 2018). These probes are quenched at acidic pH, as in late endosomes, and become fluorescent when endosomes fuse with the plasma membrane. In VPS41 KO cells, we observed multiple co-fusion events of CD63 and APP where both fluorescent signals spatiotemporally correlated, indicating that they are released from the same endosome, resulting in the release of exosomes (**Fig. 8F-H, Video S4**).

Together, these data show that depletion of the HOPS complex promotes the secretion of EVs containing APP-CTFs, but not flAPP. This increase in secretion was most prominent in VPS41 KO cells.

## Discussion

Defects in the HOPS complex have been linked to neurological disorders, including AD (Harrington et al., 2012; Zhang et al., 2016; Griffin et al., 2018; Welle et al., 2021; Monfrini et al., 2021; Polanco et al., 2022;). However, the cellular cascades linking HOPS impairment to neurodegeneration remain poorly understood. Additionally, it is unclear whether the reduced expression patterns of HOPS subunits during AD could potentially play a role in moderating disease progression (Dharshini et al., 2019; Polanco et al., 2022). Here, we studied the role of HOPS in APP trafficking and processing. We find that HOPS depletion impairs APP recycling from endosomes to the TGN, which is likely due to a failure to enter retromer-positive recycling tubules (**Fig. 8I**). Consequently, APP in HOPS KO cells accumulates in LAMP1-positive late endosomes, where it co-distributes with aberrantly localized BACE1, resulting in toxic APP-CTFs buildup (**Fig. 8I**). This phenotype is reminiscent of multiple models of AD, in which APP and its metabolites accumulate in LAMP1-positive compartments (Gowrishankar et al., 2015b; Kaur et al., 2016). Intriguingly, we observed increased secretion of APP-CTFs via EVs, which are formed as ILVs in late endosomes and secreted by fusion of these late endosomes, or multivesicular bodies, with the plasma membrane. The latter pathway is upregulated in HOPS KO cells, while fusion of late endosomes with lysosomes, which is HOPS dependent, is obstructed (**Fig. 8I**) (Shelke et al., 2023). Since the accumulation of APP-CTFs can damage endosomes and further promote EV release (Lauritzen et al., 2019; Bretou et al., 2024), their secretion by HOPS-impaired cells may contribute to disease propagation by compromising the endo-lysosomal system in neighboring cells that internalize EVs by endocytosis (Laulagnier et al., 2017; Sardar Sinha et al., 2018).

To study the trafficking of newly synthesized APP in the presence of absence of the HOPS complex, we applied the previously established RUSH tool, which allowed us to systematically track APP transport upon release from the ER (Boncompain et al., 2012). The trafficking of RUSH-APP from the ER to the Golgi and subsequent release were similar in WT and VPS41 KO cells. Previously, we reported a novel and independent role for the HOPS subunit VPS41 in the direct delivery of newly synthesized lysosomal membrane proteins (e.g. LAMPs, V0-ATPase, NPC1) to late compartments of the endo-lysosomal system (Sanzà et al., 2025; Pols et al., 2013; Swetha et al., 2011). Our present data in VPS41 KO cells indicate that APP is not transported via this pathway. Consistent with previous studies, we observe that biosynthetic APP enters the endo-lysosomal system in EEA1-positive early endosomes in WT cells (Toh et al., 2017). In HOPS KO cells, most EEA1-positive compartments correspond to enlarged hybrid early-to-late endosomes that also accumulate autophagic material (van der Beek et al., 2024). Still, also in these HOPS depleted cells, biosynthetic APP efficiently reaches EEA1-positive endosomes within 1h release from the ER, indicating that HOPS is not required for the fusion of biosynthetic APP carriers with EEA1-positive endosomes.

To our surprise, we found that HOPS KO prevents efficient endosome-to-TGN recycling of APP. APP recycling is regulated by the retromer complex and endosomal sorting receptor SorL1, both of which have been implicated in AD. SorL1 can bind both VPS35 and APP, thereby functioning as a moderator of endosome-to-TGN transport of APP (Fjorback et al., 2012). Loss-of-function variants of SorL1 have been identified in patients with late-onset AD, and disruption of the retromer subunits VPS35 and VPS26 has been shown to enhance amyloidogenic processing of APP in AD models (Jensen et al., 2024; Holstege et al., 2017; Raghavan et al., 2018; Small et al., 2005). How could depletion of HOPS, the complex required for lysosomal fusion, perturb APP recycling? Previously, we have shown that HOPS-depletion also blocks endosome-to-TGN recycling of CI-M6PR (van der Beek et al., 2024), resulting in its accumulation in aberrant, enlarged EEA1- and LAMP1-positive endosomes (HOPS bodies) that were not properly acidified. Since an acidic pH is essential for ligand dissociation from CI-M6PR and a pre-requisite for its recycling, the failure to recycle CI-M6PR in HOPS KO cells was ascribed to the elevated pH in HOPS bodies (Braulke & Bonifacino, 2009). In contrast to CI-M6PR, APP accumulated in LAMP1-positive/EEA1-negative late endosomes in HOPS KO cells. However, the RUSH-APP experiments demonstrated that newly synthesized APP transiently traffics through EEA1-positive endosomes before accumulating in late endosomes. Interestingly, SorL1 normally localizes to EEA1- and VPS35-positive endosomes and recycling tubules (Pietilä et al., 2019) and endosomal acidification is also critical for SorL1-mediated recycling of APP to the TGN (Mehmedbasic et al., 2015). Therefore, in HOPS KO cells, the elevated pH of EEA1-positive HOPS bodies may impair the binding of APP to SorL1, thereby disrupting its recycling and leading to the misrouting of APP to late endosomes. Because HOPS deficiency impairs endosome–lysosome fusion, APP subsequently accumulates in these compartments.

An additional interesting possibility is that HOPS depletion could generally affect the function of the retromer complex. Retromer function depends on the sequential action of RAB5 and RAB7 (Rojas et al., 2008). The RAB7 activation state, which is regulated by the RAB-GAP TBC1D5, is important for retromer association and dissociation from membranes (Rojas et al., 2008; Jia et al., 2016). Interestingly, HOPS depleted cells display delayed RAB5 to RAB7 switching, which could potentially impact RAB7 activation (van der Beek et al., 2024; Rink et al., 2005). Furthermore, a recent study indicates a potential interaction between HOPS subunit VPS39 and TBC1D5, suggesting that HOPS could regulate retromer association and dissociation with endosomes (Walia et al., 2024). Reduced recruitment of TBC1D5 may lead to RAB7 hyperactivation, impairing VPS35 dissociation from endosomes and ultimately disrupting endosome-to-TGN recycling (Walia et al., 2024; Daly et al., 2023; Xie et al., 2020). Future studies are needed to elucidate the exact interplay and dynamics of HOPS and retromer and how this is intertwined with trafficking of key cargo proteins, including APP.

We showed that HOPS disruption induces co-distribution of both APP and BACE1 in late endosomes, promoting CTF accumulation, but not Aβ production. Since lysosomes are the primary sites for degradation and HOPS depletion impairs late endosome – lysosome fusion, APP degradation was expected to be impaired in these cells. Indeed, by western blotting we found increased levels of flAPP in HeLa HOPS KO cells, most significantly upon VPS41 KO. Since we found that knockdown of VPS41 was lethal in neurons, we alternatively expressed the SARS-CoV2 accessory protein ORF3a to impair HOPS function (Miao et al., 2020; Walia et al., 2024). A benefit of this approach is that it allows for rapid disruption of HOPS functioning (∼24h) (Walia et al., 2024). A downside is that ORF3a also induces cytokine production and caspase activation (Zhang et al., 2022). However, since expression of ORF3a mutants that fail to bind VPS39 (Chen et al., 2021; Walia et al., 2024) did not deplete APP from the Golgi-region, the effects of ORF3a expression on APP localization appeared to be HOPS specific. Of note, in contrast with HeLa HOPS KO cells, we did not observe an increase in flAPP levels in ORF3a expressing neurons. Since ORF3a expression was done for a maximum of 7 days, as opposed to several passages of HeLa HOPS KO cells, it is likely that only prolonged HOPS disruption leads to accumulation of flAPP.

Since endosomes are a dominant site for amyloidogenic APP processing (Park et al., 2022), this led us to ask whether APP-positive endosomes in HOPS depleted cells also contained amyloidogenic secretases involved in APP cleavage. Interestingly, we found increased co-distribution of APP and the β-secretase BACE1 in these conditions. Normally, BACE1 is sorted from early endosomes to recycling endosomes destined for the plasma membrane and less frequently to late endosomes (Das et al., 2013; Toh et al., 2018; Tan and Gleeson, 2019b). BACE1 trafficking to recycling endosomes has been shown to be partly regulated by retromer (Toh et al., 2018; Tan and Gleeson, 2019b). Therefore, the reported defects in endosomal maturation and recycling upon HOPS disruption (van der Beek et al., 2024) may also prevent the sorting of BACE1 into these recycling endosomes, with BACE1 ending up in the same late endosome compartments as APP. Consistent with the observed increase in co-distribution of APP and BACE1, we also observed increased levels of APP-CTFs by western blot. Furthermore, fluorescent imaging showed segregation of dual N- and C-terminal APP tags in HeLa KO cells, as well as in ORF3a-expressing neurons, indicating increased processing of APP. By comparing processing of APP_ice_ and APP_WT_ in HOPS KO cells, we found that this increase in processing was partly caused by BACE1 cleavage and that both C83 as C99 accumulate in HOPS KO cells. Surprisingly, recent work demonstrated that both C83 and C99 cause a decrease in endo-lysosomal Ca^2+^ and an increase in endosomal cholesterol, emphasizing that efficient clearance of both APP-CTFs is required for proper endo-lysosomal homeostasis (Bretou et al., 2024). We conclude that HOPS KO also impairs the sorting of BACE1, resulting in increased co-distribution of APP an BACE1 in late endosomes, thereby promoting the generation of APP-CTFs.

In neurons, we found that the endosomal build-up of flAPP and APP-CTFs was confined to the somatodendritic domain, suggesting a role for lysosomes within this domain in degrading APP fragments. Additionally, these APP compartments were mostly LAMP1-positive and stationary. In healthy neurons, endo-lysosomal maturation depends on retrograde transport to the perinuclear area, where most catalytic lysosomes are concentrated (Ferguson, 2018; Yap et al., 2022, 2018). Impaired HOPS function may disrupt this maturation process and hinder transport and fusion of late endosomes with lysosomes, resulting in defective degradation and localized accumulation of APP-CTFs in the somatodendritic domain. Currently, our understanding of the regulation of endo-lysosomal transport within dendrites remains limited, partly due to their complex and mixed microtubule organization (Kapitein and Hoogenraad, 2011). Interestingly, recent findings indicate that RAB7 activation is essential for transport of specific dendritic proteins, such as Nsg1 and Nsg2 (Yap et al., 2018). Since HOPS impaired cells display delayed RAB5 to RAB7 conversion and RAB7 hyperactivation (Walia et al., 2024; van der Beek et al., 2024; Rink et al., 2005), it is plausible that these disruptions affect dendritic lysosome dynamics. However, how RAB7 hyperactivation specifically influences dendritic transport of lysosomes remains an open question.

We expected that APP-CTF accumulation upon HOPS disruption would induce Aβ production in neurons, as the γ-secretase subunit PSEN2 normally localizes to late endosomes and lysosomes (Sannerud et al., 2016). However, HOPS disruption by ORF3a overexpression did not increase Aβ42 levels. Of note, HeLa HOPS KO cells also contain a population of enzymatically active lysosomes, though these are poorly reached by cargo’s destined for degradation (van der Welle et al., 2021; van der Beek et al., 2024). Upon HOPS disruption, delivery of APP-CTFs to PSEN2-positive endo-lysosomal compartments is delayed, which explains the similar Aβ42 levels detected in control and ORF3a expressing cells. In addition, ORF3a-mediated disruption of HOPS over 7 days may not be sufficient to induce the accumulation of APP-CTFs and PSEN2 within the same endosomes at levels high enough to produce detectable amounts of Aβ42. Interestingly, APP-CTF associated neuronal impairments actually precede Aβ formation in 3xTg-AD mice (APP Swedish mutation, MAPT P301L and PSEN1 M146V), suggesting that only prolonged APP-CTF accumulation and defective lysosomal clearance lead to increased Aβ production in neurons (Lauritzen et al., 2012).

Intriguingly, we found that APP-CTFs accumulated in late endosomes upon HOPS disruption were subsequently secreted in EVs, which could potentially have consequences for disease propagation. Previous work has demonstrated that defects in endo-lysosomal maturation drastically increases EV release, however, it was yet unclear to what extend these EVs contain flAPP and/or its metabolites (Shelke et al., 2023). Our findings suggests that depletion of HOPS, which decreases fusion of late endosomes with lysosomes, induces secretion of APP-CTFs by increasing fusion of late endosomes with the plasma membrane, resulting in the release of APP-CTF-containing EVs. KO of VPS41 resulted in a more drastic release of EVs than KO of other subunits, consistent with higher levels of CTFs in this cell line. This drastic phenotype can possibly be attributed to the unique role of VPS41 in lysosomal biogenesis (Pols et al., 2013; Sanzà et al., 2025), leading to more pronounced lysosomal dysfunction. Secretion of APP-derived fragments may add to AD progression by propagation to other neurons (Xiao et al., 2017). APP-CTFs can be secreted in EVs and endocytosed by neighboring cells (Perez-Gonzalez et al., 2012; Laulagnier et al., 2017 Lauritzen et al., 2019; Cone et al., 2020). APP-CTFs can be further processed by the recipient cells, indicating a possible mechanism for disease spreading (Laulagnier et al., 2017). For future research, HOPS and in particular its subunit VPS41 could provide promising targets for disease treatment. Overexpression of VPS39, VPS41 and the HOPS effector ARL8B have already been shown to reduce Aβ-induced neurodegeneration in *C. elegans*, indicating a possible role in neuroprotection (Griffin et al., 2018). Optimization of similar approaches in more complex animal models of AD could provide a first step towards the development of novel therapeutic strategies.

## Materials and methods

### Animals

All experiments involving animals were approved by the DEC Dutch Animal Experiments Committee (Dier Experimenten Commissie), performed in line with institutional guidelines of University Utrecht, and conducted in agreement with Dutch law (Wet op de Dierproeven, 1996) and European regulations (Directive 2010/63/EU). The animal protocol has been evaluated and approved by the national CCD authority (license AVD10800202216383). Female pregnant Wistar rats were obtained from Janvier, and embryos (both genders) at embryonic (E)18 stage of development were used for primary cultures of hippocampal and cortical neurons. The animals, pregnant females and embryos have not been involved in previous procedures.

### Primary neuron culture and transfection

Hippocampi or cortices from embryonic day 18 rat brains were dissected and dissociated with trypsin for 15 min and plated on coverslips coated with poly-L-Lysine (37.5µg/mL) and laminin (1.25µg/mL) at a density of 100,000/well (12-well plates) for imaging (live-cell and confocal microscopy) or 1,000,000/well (6-well plates) for biochemical assays (Western Blot and ELISA). The day of neuron plating corresponds to day-in-vitro 0 (DIV0). Neurobasal medium (NB) supplemented with 2% B27 (GIBCO), 0.5 mM glutamine (GIBCO), 15.6 µM glutamate (Sigma), and 1% penicillin/streptomycin (GIBCO) was used to maintain neurons incubated under controlled temperature and CO2 (37 °C, 5% CO_2_).

Hippocampal neurons were transfected using Lipofectamine 2000 (Invitrogen). Briefly, DNA was mixed with 3.3 μL of Lipofectamine 2000 in 200 μL NB medium containing no supplements and incubated for 20 minutes at room temperature. DNA mix was subsequently added to neurons and incubated for 1h (37 °C, 5% CO2). Next, neurons were washed with NB and transferred to half of their original medium and half fresh supplemented NB at 37 °C in 5% CO2 until fixation or imaging at different days in vitro (DIV) as indicated in figure legends.

### Culture of cell lines and transfection

HeLa parental cells were obtained from DSMZ (ACC 57). HeLa HOPS knock-out cells (VPS18 KO, VPS39 KO, VPS41 KO) were generated as previously described (van der Welle et al., 2021). HEK293T cells were obtained from ATCC and were a kind gift from Harold MacGillavry. Cell lines were maintained in DMEM (HPSTA; Capricorn) supplemented with 10% FBS and 1% penicillin and streptomycin. Cells were kept at 37°C and 5% CO_2_. HeLa and HEK293T cells were passaged twice or thrice per week respectively after reaching 70-90% confluence. Cells were transfected after reaching 70% confluence. For electron microscopy, HeLa cells were transfected with Effectene and plasmid mix ON. In brief, 0.6 ug DNA per well (6 well plate) was mixed with 4.8 µL Enhancer in EC buffer and incubated 5 minutes at room temperature. Next, 12 µL Effectene was added to the mixture followed by 10-minute incubation. DNA mix was added to cells and incubated ON (37 °C, 5% CO_2_) until fixation. For live-cell imaging and confocal microscopy, cells were transfected with Lipofectamine 2000. In brief, DNA was mixed with 2 μL of Lipofectamine 2000 in 25 μl optiMEM (GIBCO) containing no supplements and incubated for 20 minutes at room temperature. DNA mix was subsequently added to cells and incubated for 2-3h (37 °C, 5% CO_2_). Following incubation with DNA mix, medium was replaced to supplemented DMEM and incubated ON (37 °C, 5% CO_2_) until fixation or imaging.

### Exosome collection

HeLa WT and HOPS KO cells were plated in 15cm dishes and cultured in supplemented DMEM ON (37 °C, 5% CO_2_). 24h after plating, supplemented DMEM was changed to EV free media. EV free media was prepared by centrifuging FBS at 100,000x g overnight. The supernatant of the centrifuged FBS was filtered through a 0.2 μm filter and added to the media (DMEM (HPSTA; Capricorn) supplemented with 10% FBS and 1% penicillin and streptomycin).

Media was collected from plates 72h after addition of EV depleted media. Cells were lysed according to standard procedure (*Cell lysis and Western Blot*). Remaining cells, cell debris and apoptotic bodies were removed from the collected conditioned media by consecutive centrifugation at 200g for 5 min, 500g for 5 min, followed by 2000g for 20 min. All further steps were conducted at 4 °C to avoid degradation of material. Supernatant was additionally concentrated using a 10 kDa centrifugal filter (Amicon® Ultra-4). Remaining volume was fractionated based on size exclusion tomography (IZON), resulting in 13 different fractions (1-3 EV fraction, 4-13 soluble fraction). Both pooled fractions were concentrated to a total final volume of 500 μL. For Western blot analyses, EV volume used was normalized to protein concentration of cell lysates to correct for variations in confluency between cell lines.

### Lentivirus packaging and transduction in neurons

HEK293T cells were transfected at 70% confluence with packaging vector psPAX2 and envelope vector pMD2.G with a ratio of 3:2:1 (mCherry / ORF3a-mCherry) or 4:2:1 (HaloTag-APP-mNeonGreen) using PEI Max. Transfection mix was prepared in optiMEM (GIBCO) and done according to standard protocols with a 3:1 PEI Max:DNA ratio and followed by 20-minute incubation at room temperature. DNA mix was subsequently added to HEK293T cells and incubated for approximately 6h (37 °C, 5% CO_2_). Medium was removed after 6h and changed to NB supplemented with 0.5 mM glutamine. Supernatant from packaging HEK293T cells was collected 48 to 72h after transfection and filtered through a 0.45 µm filter to remove remaining cell debris. Supernatant was further concentrated using an Ultra Centrifugal Filter 100 kDa MWCO (AMICON) at 4°C 1,000 g for 15 to 30 min to a final volume of ±100 μL. Concentrated virus was stored at 4 °C for short term storage (max. 2 weeks) or −80 °C long term storage until use. Day of lentiviral transduction is indicated in figure legends. 1-2 μL (12-well plate) or 10-20 μL (6-well plate) of concentrated virus was used per construct. Respective days of lentiviral transduction are indicated in figure legends.

### DNA and shRNA constructs

The following constructs were used in this study:

pEGFP-n1-APP was a gift from Zita Balklava & Thomas Wassmer (Addgene plasmid #69924) (Currinn et al., 2016), HaloTag-CLCa was a gift from Harold MacGillavry (Catsburg et al., 2022), FUGW was a gift from David Baltimore (Addgene plasmid #14883) (Lois et al., 2002), psPAX_2_ and pMD2.G were gifts from Didier Trono (Addgene plasmids #12260 and #12259), pmScarlet-i_C1 was a gift from Dorus Gadella (Addgene plasmid #85044) (Bindels et al., 2016), H2B-NeonGreen-IRESpuro2 was a gift from Daniel Gerlich (Addgene plasmid #183745) (Cuylen et al., 2016), pCMV-Sport6-CD63-pHluorin was a gift from DM Pegtel (Addgene plasmid #130901) (Droogers et al., 2022), ORF3a-mCherry was a gift from Bruno Antonny (Addgene plasmid #165138) (Miserey-Lenkei et al., 2021), and Str-KDEL_SBP-EGFP-Ecadherin was a gift from Franck Perez (Addgene plasmid #65286) (Boncompain et al., 2012).

The following plasmids were generated for this study (all primers used in this study can be found in **Table 1**):

**Table 1:**
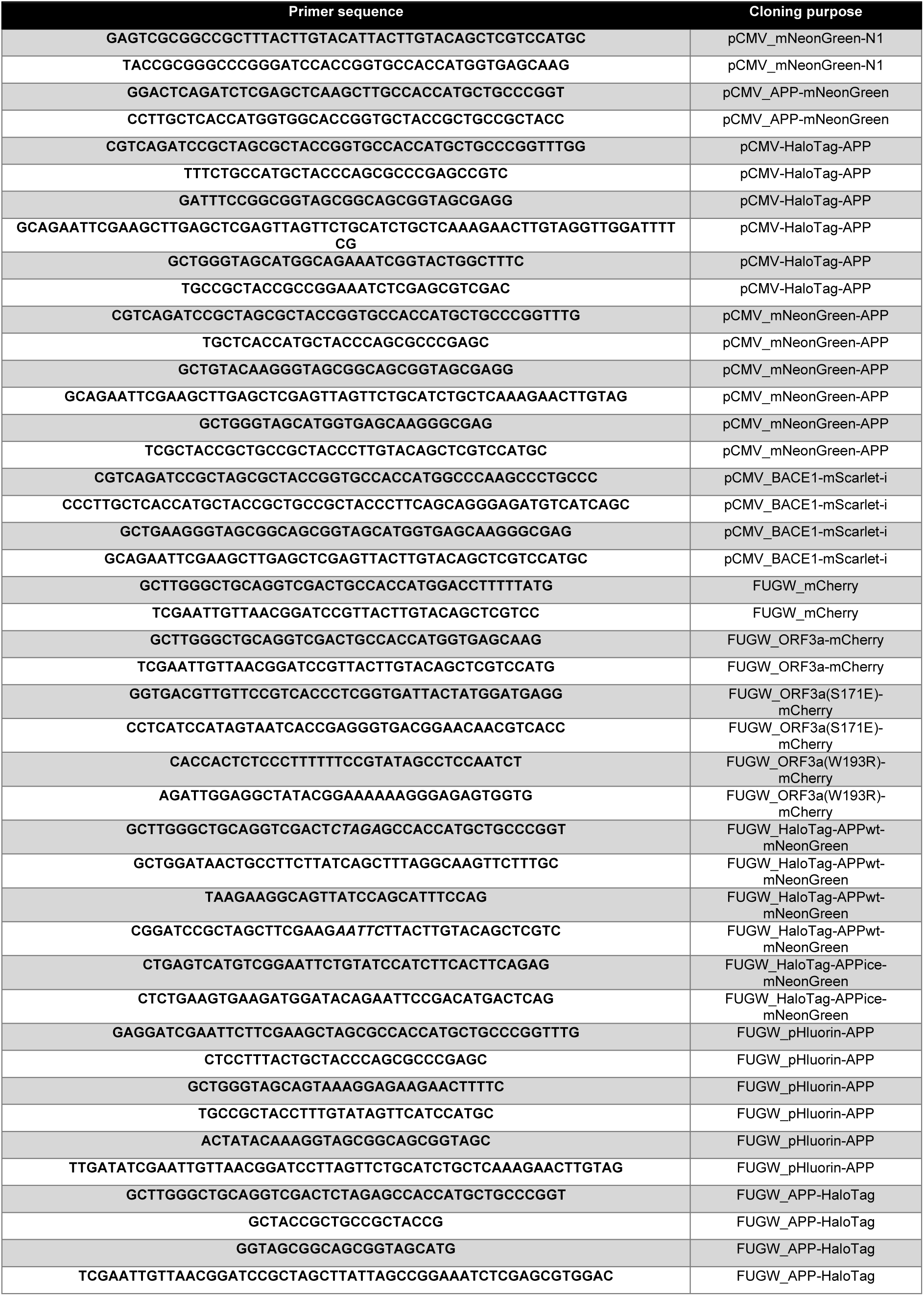
Primer table.

The following plasmids were generated for this study:

To generate APP-mNeonGreen, we first generated mNeonGreen-N1, by removing mEGFP from pEGFP-N1 with AgeI and BsrGI. mNeonGreen was amplified from H2B-mNeonGreen (Cuylen et al., 2016) and inserted in the vector. mNeonGreen-N1 was subsequently digested with AgeI and HindIII. APP was amplified from pEGFP-N1-APP (Addgene plasmid #69924) (Currinn et al., 2016) and inserted in open mNeonGreen-N1. For the cloning of pCMV HaloTag-APP and pCMV mNeonGreen-APP, pEGFP-C1 was digested using XhoI and AgeI. HaloTag was amplified from HaloTag-CLCa (Catsburg et al., 2022), mNeonGreen was amplified from H2B-mNeonGreen and APP was amplified from pEGFP-N1-APP (Addgene plasmid #69924) (Currinn et al., 2016). All fragments were inserted using Gibson Assembly. For the cloning of FUGW APP-HaloTag, FUGW (Addgene plasmid #14883) (Lois et al., 2002) was restricted using NheI and EcoRI. APP was amplified using PCR from pCMV APP-mNeonGreen and HaloTag was amplified from BACE1-HaloTag (from Farías lab).

For the generation of SBP-mNeonGreen-APP, FUGW-RUSH (Li et al., 2024) was restricted using BamHI and EcoRI. SBP was amplified from Str-KDEL_SBP-EGFP-Ecadherin (Addgene plasmid #65286) (Boncompain et al., 2012) and APP signal peptide and mNeonGreen-APP from pCMV mNeonGreen-APP. All fragments were inserted using Gibson Assembly.

For HaloTag-APP-mNeonGreen constructs, FUGW (Addgene plasmid #14883) (Lois et al., 2002) digestion was done using EcoRI-HF and XbaI. HaloTag-APP_(half)_ and _(half)_APP-mNeonGreen were amplified from previous generated vectors and inserted in open FUGW. The Icelandic mutation was introduced in this construct by PCR of HaloTag-APP-mNeonGreen using primers containing respective mutation. Gibson Assembly was used for all ligation reactions.

To generate C-terminally tagged BACE1-mScarlet, pmScarlet-i_C1 (Addgene plasmid #85044) (Bindels et al., 2016) was digested using XhoI and AgeI, resulting in the removal of mScarlet from the plasmid. BACE1 was amplified from rat cDNA acquired from I.M.A.G.E. consortium and mScarlet was amplified from pmScarlet-i_C1. Both fragments were inserted in open vector using Gibson Assembly.

For the generation of FUGW_mCherry and FUGW_ORF3a-mCherry constructs, FUGW vector (Addgene plasmid #14883) (Lois et al., 2002) was digested using NheI and XbaI. mCherry was amplified from pGW2-mCherry and ORF3a from ORF3a-mCherry (Addgene plasmid #165138) (Kapitein et al., 2010b). Fragments were inserted using Gibson Assembly. Mutations in FUGW ORF3a-mCherry (S171E and W193R) were generated by using the QuickChange II XL Site-Directed Mutagenesis Kit.

The following sequences for rat-shRNAs inserted to pLKO.1-puro and/or pSuper were used in this study: shScramble (5’-ACAATAGCTTACTACCAAT-3’), shVPS41#1 (5’-CACTCGTTCAATGACGTGT-3’), shVPS41#2 (5’-ATTGACAAACCACCATTTA-3’), shVPS41#3 (5’-CCACTCATCTATGAAATGA-3’), shVPS41#4 (5’-GCTTTGATGGCAGCTGAAA-3’). Knockdown efficiencies were validated by comparing VPS41 immunolabel intensity in empty pSuper / shVps41 transfected cells co-transfected with pEGFP-N1 (**Fig. S3A-B**).

### Reagents and antibodies

For antibodies used in this study, see **Table 2**. Other reagents used in this study were: Effectene (QIAGEN, #301425), Lipofectamine 2000 (Invitrogen, #1639722), PEI (Polysciences, #19850), SiR-lysosome kit (Spirochrome, #SC012), LysoTracker™ Red DND-99 (Thermo Fisher Scientific, #L7528), biotin (Sigma-Aldrich, #B4501), Gibson Assembly Mix (New England Biolabs, #E2611), DAPT γ-secretase inhibitor (Abcam, #ab120633), Human/Rat β Amyloid 42 ELISA Kit Wako High Sensitive (FUJIFILM, #292-64501), JFX650 was kindly provided by the Lavis Lab (Janelia).

**Table 2:** Antibody table.

| Name | Host | Catalog number | Company | Dilution |
| --- | --- | --- | --- | --- |
| PRIMARY ANTIBODIES |  |  |  |  |
| APP | Rabbit | ab32136 | Abcam | 1:200 (IF) 1:1000 (WB) |
| LAMP1 | Mouse | 555798 | BD Pharmingen | 1:500 (IF) |
| EEA1 | Mouse | 610456 | BD Biosciences | 1:100 (IF) |
| EEA1 | Rabbit | C45B10 | Cell Signaling | 1:200 (IF) |
| GM130 | Mouse | 610823 | BD Biosciences | 1:500 (IF) |
| GM130 | Rabbit | ab52649 | Abcam | 1:800 (IF) |
| VPS41 | Mouse | SC-377271 | Santa Cruz | 1:250 (IF) |
| VPS35 | Goat | ab10099 | Abcam | 1:200 (IF) |
| VPS35 | Mouse | sc-374372 | Santa Cruz | 1:1000 (WB) |
| SNX1 | Rabbit |  | Gift from Jez Carlton | 1:250 (IF) |
| CI-MPR | Rabbit |  | Gift from Stuart Kornfeld | 1:500 (IF) |
| BACE1 | Rabbit | 5606S | Cell Signaling | 1:1000 (WB) |
| PSEN2 | Rabbit | ab51249 | Abcam | 1:20000 (WB) |
| Alpha-tubulin | Mouse | T5168 | Sigma | 1:20000 (WB) |
| GAPDH | Rabbit | G9545 | Sigma | 1:5000 (WB) |
| HaloTag | Rabbit | G9281 | Promega | 1:30 (EM) |
| TRIM46 | Guinea Pig | 377005 | Synaptic Systems | 1:1000 (IF) |
| CD63 | Mouse | 556019 | BD Biosciences | 1:750 (WB) |
| Calnexin | Rabbit | ab22595 | Abcam | 1:1000 (WB) |
| SECONDARY ANTIBODIES |  |  |  |  |
| Anti-mouse HRP | Rabbit | P0260 | Agilent | 1:10000 (WB) |
| Anti-rabbit HRP | Swine | P0399 | Agilent | 1:10000 (WB) |
| anti-Mouse IRDye 680 | Donkey | 926-68020 | LICORbio | 1:20000 (WB) |
| anti-Rabbit IRDye 680 | Donkey | 926-68021 | LICORbio | 1:20000 (WB) |
| anti-Mouse IRDye 800 | Donkey | 926-32210 | LICORbio | 1:20000 (WB) |
| anti-Rabbit IRDye 800 | Donkey | 926-32211 | LICORbio | 1:20000 (WB) |
| Anti-Guinea Pig CF405S | Donkey | SAB4600230 | Sigma | 1:500 (IF) |
| Anti-Mouse Alexa 488 | Goat | A11029 | Life Technologies | 1:1000 (IF) |
| Anti-Rabbit Alexa 488 | Goat | A11034 | Life Technologies | 1:1000 (IF) |
| Anti-Mouse Alexa 647 | Goat | A21236 | Life Technologies | 1:1000 (IF) |
| Anti-Rabbit Alexa 647 | Goat | A21245 | Life Technologies | 1:1000 (IF) |
| Anti-goat Alexa 647 | Donkey | A21447 | Life Technologies | 1:1000 (IF) |

### Cell lysis and Western Blot

HeLa cells and primary rat cortical neurons were culture in 6-well plates and washed two times in ice cold PBS before scraping. For the improved detection of APP-CTFs, cells were incubated with 25 µM γ-secretase inhibitor DAPT (Abcam) or DMSO control for 20h prior to cell lysis. Cells were scraped and lysed in RIPA buffer (150mM NaCl, 50mM Tris-HCl pH7.4, 0.1% SDS, 0.5% sodium-deoxycholate, 1% Triton X-100 and 1x protease inhibitor (Roche)) followed by incubation for 15 min at 4 °C. Lysates were cleared by centrifuging at 16,000 x g 4°C after which the pellet was discarded. Protein concentration was measured using the Pierce™ BCA Protein Assay Kit (Thermo Fisher Scientific) in order to equalize protein loading on gels. 20 µg of protein was used for all experiments, unless indicated differently in figure legend. Lysates were resolved by SDS-PAGE on a 12% Bis-Acrylamide (Bio-Rad) gel and transferred to a PVDF membrane (Bio-Rad). These membranes were blocked for 1h at room temperature in TBS-T containing 3% BSA. Primary antibodies were diluted in blocking buffer and incubated ON at 4 °C. After three washes with TBS-T, membranes were incubated with HRP or IRDye secondary antibody in blocking buffer for 1h at room temperature and washed three times with TBS-T prior to imaging. For samples incubated with HRP secondary antibody: membranes were incubated in Clarity Western ECL Substrate (Bio-Rad) and developed using ImageQuant 800 (AMERSHAM). For samples incubated with IRDye secondary antibody: membranes were developed on an Odyssey CLx imaging system (LICOR) with Image Studio version 5.2.

### ELISA measurement of Aβ42

Cortical neurons were lysed using RIPA buffer on DIV15 as described in previous section (*Cell lysis and Western Blot*). For testing of cellular Aβ42, the Human/Rat β Amyloid 42 ELISA Kit Wako High Sensitive (FUJIFILM) was used. Standard curves were generated during every set of experiments and all samples were measured in duplicate. 20 µg protein was used for all different measurements.

### Immuno-Electron Microscopy (I-EM)

HeLa WT and HOPS KO cells were grown in 6cm dishes and transfected with HaloTag-APP or APP-HaloTag construct as described in previous section (*Culture of cell lines and transfection*). Cells were fixed the day after transfection by adding 2% FA, 0.2% GA in 0.1 M phosphate buffer (pH 7.4) to an equal volume of medium for 5 min at room temperature. Afterwards, fixative was refreshed and cells were incubated for 2h. Cells were stored in 1% formaldehyde at 4°C until further processing. Then, cells were washed with PBS/0.05 M glycine, scraped in 1% gelatin/PBS, pelleted in 12% gelatin/PBS, solidified on ice, and cut into small blocks. The blocks were infiltrated in 2.3 M sucrose overnight at 4ᵒC, mounted on aluminum pins, and frozen in liquid nitrogen (Slot and Geuze, 2007). Ultrathin sections were prepared using an ultra-cryomicrotome (Leica). To pick up ultrathin sections, a 1:1 mixture of 2.3 M and 1.8% wt/vol methylcellulose was used. Ultrathin sections were immuno-gold labeled and stained as described (Slot and Geuze, 2007). Protein A-conjugated colloidal gold particles were made in house (Cell Microscopy Core, UMC Utrecht).

### Live-cell imaging

For live-cell imaging of HeLa cells and primary rat hippocampal neurons, cells were imaged on a Nikon Eclipse Ti or Nikon Eclipse Ti2 equipped with an incubator chamber (INUG2-ZILCS0H2; Tokai Hit) on a motorized stage (ASI), a Nikon Plan Apo VC 100x N.A. 1.40 oil objective, and a spinning disk confocal scanner unit (CSU-X1-A1; Yokogawa). Illumination was performed using a CoolLED pE4000 (CoolLED) LED device and ET-EGFP (49002; Chroma), ET-mCherry (49008; Chroma), ET-CY5 (49006; Chroma), and ET-CY5.5 (49022; Chroma) filters. All images were acquired with a Coolsnap HQ2 CCD camera (Photometrics). The microscope was controlled using µManager software (Edelstein et al., 2014)(Edelstein et al., 2014)(Edelstein et al., 2014). Coverslips were mounted in a metal ring and cells were supplemented in their original medium. Cells were imaged in an incubation chamber that maintains optimal temperature and CO_2_ (37 °C and 5% CO_2_). For dual color imaging, sequential imaging was used with a maximum exposure of 0.3 seconds per channel. For imaging of constructs containing HaloTag, cells were incubated with JFX650 (100 nM) in NB without supplements prior to imaging (37 °C and 5% CO_2_). For the RUSH assay using live cells, biotin was added on stage (100 µM) for synchronized release of APP. Total time and intervals of imaging acquisition for each experiment are indicated in figure legends.

For TIRF microscopy experiments, cells were imaged 1 day post transfection with CD63-pHmScarlet and pHluorin-APP on a TIRF microscope (Nikon Eclipse Ti with Perfect Focus System, Nikon Apo TIRF 100x N.A. 1.49 oil immersion objective, Prime BSI sCMOS camera with 14.71 pixels/μm resolution, Tokai Hit INUBG2E-ZILCS stage top incubator, MS-2000-XY motorized stage (ASI), p3E-300 illuminator (CoolLED), LB10-3 filter wheel (ASI), MetaMorph 7.7 software). Videos (total duration 3 minutes - 0.3 seconds per frame) were acquired for HeLa VPS41 KO cells.

### Immunofluorescence staining and imaging

Medium was completely removed from the well and cells were incubated at with prewarmed (37 °C) 4% paraformaldehyde supplemented with 4% sucrose in PBS for 15 min for fixation. Cells were permeabilized with 0.2% Triton X-100 in PBS supplemented with calcium and magnesium (PBS-CM) for 15 min, followed by blocking with 0.2% porcine gelatin in PBS-CM for 1h at room temperature. Cells were incubated ON with primary antibody in blocking solution at 4 °C. Incubation with secondary antibody took place for 1h at room temperature. Cells were washed three times with PBS-CM in between the above-described steps. Coverslips were mounted in Fluoromount-G Mounting Medium (ThermoFisher Scientific). Cells with similar expression levels of transfected constructs were imaged on confocal microscopes. Imaging took place on a confocal laser-scanning microscope (LSM700, with Zen imaging software (Zeiss) version 8.1.7.484) equipped with Plan-Apochromat ×63 NA 1.40 oil DIC and EC Plan-Neofluor ×40 NA1.30 Oil DIC objectives.

### Labelling of late endosomes and lysosomes using SirLyso and LysoTracker probes

For the labeling of lysosomes and late endosomes, we used SirLyso and LysoTracker. For SirLyso, a dilution of 1:500 was applied for 30 min at standard culturing conditions (37 °C and 5% CO_2_). For LysoTracker a dilution of 1:20.000 was applied for 5 min at standard culturing conditions (37 °C and 5% CO_2_). After incubation, cells were washed one time in warm PBS prior to fixation according to standard procedures.

### Image and data analysis

#### Particle and colocalization analysis

The ComDet Plug-In (Eugene Katrukha, Cell Biology, Utrecht University) for ImageJ/FIJI was used for particle detection, particle size quantification and particle number detraction (**Fig. 1C**). Additionally, colocalization of indicated compartments in HeLa cells was measured by applying the same Plug-In (**Fig. 2B, 2F, 2H, 3H**, **4C, 4E**, **5C, 5I**, **7H, S1B, S1D**). For measuring the colocalization between the C- and N-terminus of HaloTag-APP-mNG in neurons, we applied the ImageJ/FIJI plugin JACoP to obtain values for Pearson Correlation (**Fig. 7E**). Values were measured in the soma of neurons.

#### Polarity index measurement

We used a previously established method to investigate the polarized distribution of proteins, called the polarity index (PI) (Kapitein et al., 2010a) (**Fig. 6G**). Segmented lines were drawn along three dendrites using ImageJ and one portion of the axon of ∼30 μm (excluded the axon initial segment) in each image. The mean intensity of the three dendrites were subsequently averaged. Afterwards, the polarity index was measured using the following formula: PI = (I_d_ − I_a_)/(I_d_ + I_a_). I_d_ indicates the mean intensity of the three dendrites and I_a_ indicates the mean intensity of the axon. PI < 0 indicates axonal distribution, PI > 0 indicates dendritic distribution and PI = 0 stands for non-polarized distribution where I_d_ = I_a_.

#### Western blot analysis

Band intensities were measured in ImageJ/FIJI using the *Analyze>Gels* option (**Fig. 5E, 5F, 7B, 7C**, **8C, 8D, S4C, S4E**). Extracted values were divided against loading control to correct for differences in total protein levels. Fold change was subsequently calculated by dividing values against control value of each replicate.

#### Golgi intensity of APP

To measure APP intensity (mean grey value) in the Golgi region, we generated a mask of the GM130 label using ImageJ/FIJI (**Fig. 1E, 3F, 6B**). Additionally, to correct for differential expression levels we measured whole cell intensity (mean grey value) and used this for the normalization to correct for differential expression levels. In neurons, we applied a similar approach, but used the intensity measurement (mean grey value) in the soma for normalization (**Fig. 6E**).

#### Kymograph analysis

Kymographs were generated from live cell videos in ImageJ. Dendrites were identified based on morphology and segmented lines (30 μm) were drawn along their tracks and subsequently straightened. Straightened dendrites were then re-sliced followed by z-projection to obtain kymograph. Anterograde movements (towards dendrite tip) were oriented from left to right in all kymographs (**Fig. 6H, S3E**).

### Statistical analysis

Data processing and statistical analysis were performed using Excel and GraphPad Prism (GraphPad Software 8.0.2). Data normality was checked prior to statistical testing using D’Agostino–Pearson omnibus test. Unpaired t test, Mann-Whitney test, Welch’s t test, Ordinary one-way ANOVA followed by Sidak correction, Kruskal-Wallis test followed by Dunn correction, Ordinary two-way ANOVA followed by Tukey correction, were performed for statistical analyses and are indicated in figure legends. The statistical test performed, P value, number of cells and independent experiments are indicated in figures and corresponding legends.

## Data availability

Raw data of all quantifications and uncropped Western blots are available in the Data File. Any other data reported in this paper will be shared by the lead contact upon reasonable request.

## Acknowledgments

We thank Volker Haucke (Leibniz-FMP, DE), David Gershlick (Cambridge Institute for Medical Research, UK), Max Koppers (Vrije Universiteit Amsterdam, NL), and Maarten Bebelman (Utrecht University, NL) for their valuable suggestions and help with experimental design. We thank the Klumperman lab and Farías lab and in particular Paolo Sanza and Jan van der Beek for fruitful discussions and sharing HeLa HOPS KO lines.

## CRediT authorship contribution statement

**D.B.H. Draper:** conceptualization, formal analysis, investigation, methodology, visualization, writing – original draft. **A.E. George:** formal analysis, investigation, methodology. **T. Veenendaal:** investigation. **S. van Dijk:** investigation. **F.J. Verweij:** methodology, supervision, writing – review & editing. **J. Klumperman:** conceptualization, funding acquisition, methodology, project administration, supervision, writing – review & editing. **G.G. Farías:** conceptualization, funding acquisition, methodology, project administration, supervision, writing – review & editing.

## Declaration of generative AI and AI-assisted technologies in the writing process

During the preparation of this work the authors used ChatGPT in order to improve the readability of this manuscript. After using this tool, the authors reviewed and edited the content as needed and take full responsibility for the content of the publication.

---

**Figure S1.**
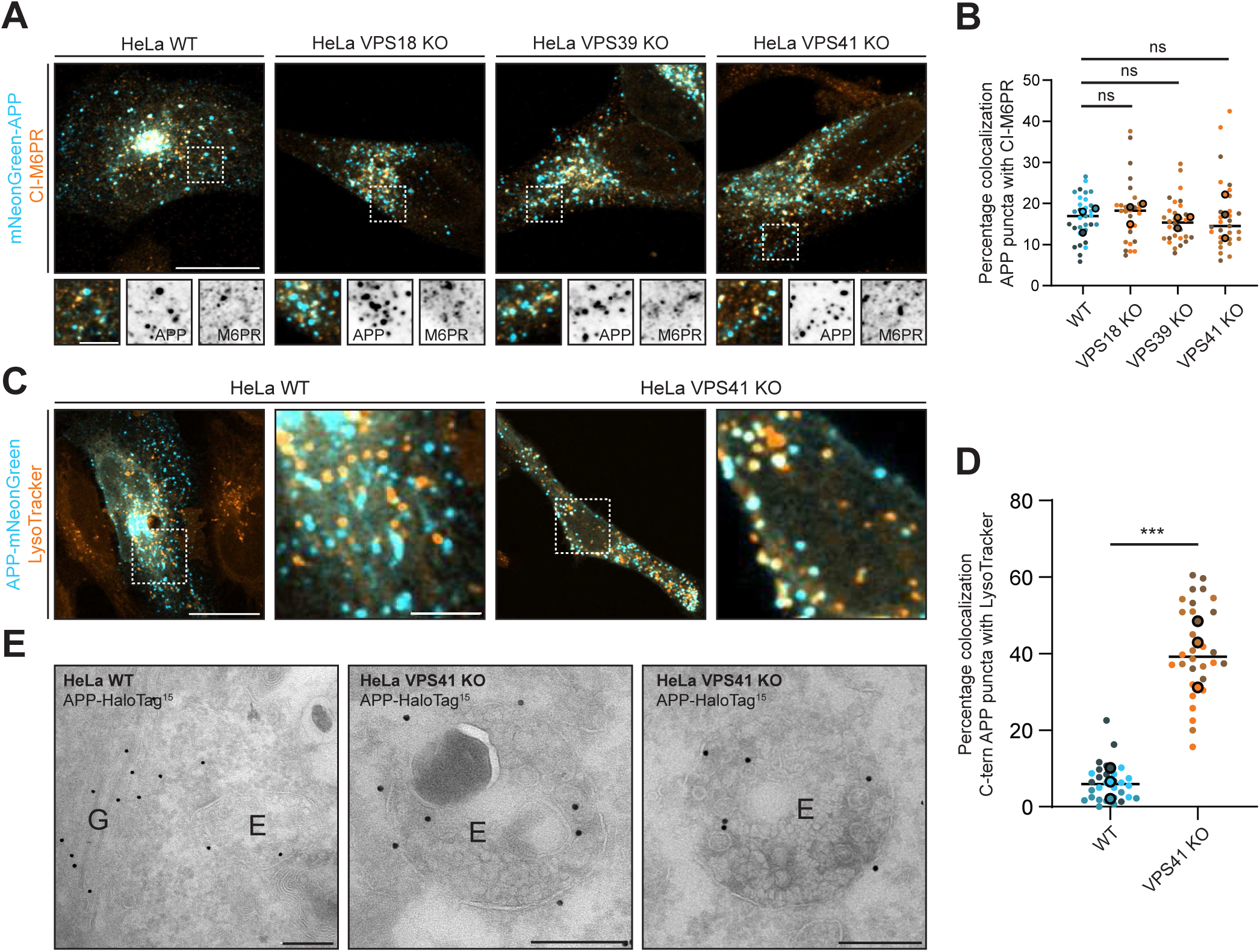
APP and CI-M6PR are distributed in distinct endosomal populations in both HeLa WT and HOPS KO cells. **A)** Representative images of HeLa WT and HOPS KO cells expressing mNeonGreen-APP (blue) and stained for endogenous CI-M6PR (orange). **B)** Quantifications of the percentage of APP puncta positive for CI-M6PR. **C)** Representative images of HeLa WT and VPS41 KO cells expressing APP-mNeonGreen and labelled with LysoTracker probe (orange). **D)** Quantification of the percentage of C-terminal APP puncta positive for LysoTracker. **E)** Immuno-EM of HeLa WT and VPS41 KO cells expressing APP-HaloTag (15 nm gold particles) G = Golgi, E = Endosome. All scale bars indicate a region of 20 µm (overview), 5 µm (zooms) or 200 nm (EM). Dots in all dot plots indicate individual cells and lines indicate median value. Mean values of each replicate are indicated in dot blots by bold circles. Kruskal-Wallis test followed by Dunn’s correction for multiple comparisons was used in (B). Unpaired t test with Welch’s correction was used in (D). ns = not significant, *** (P < 0.001).

**Figure S2.**
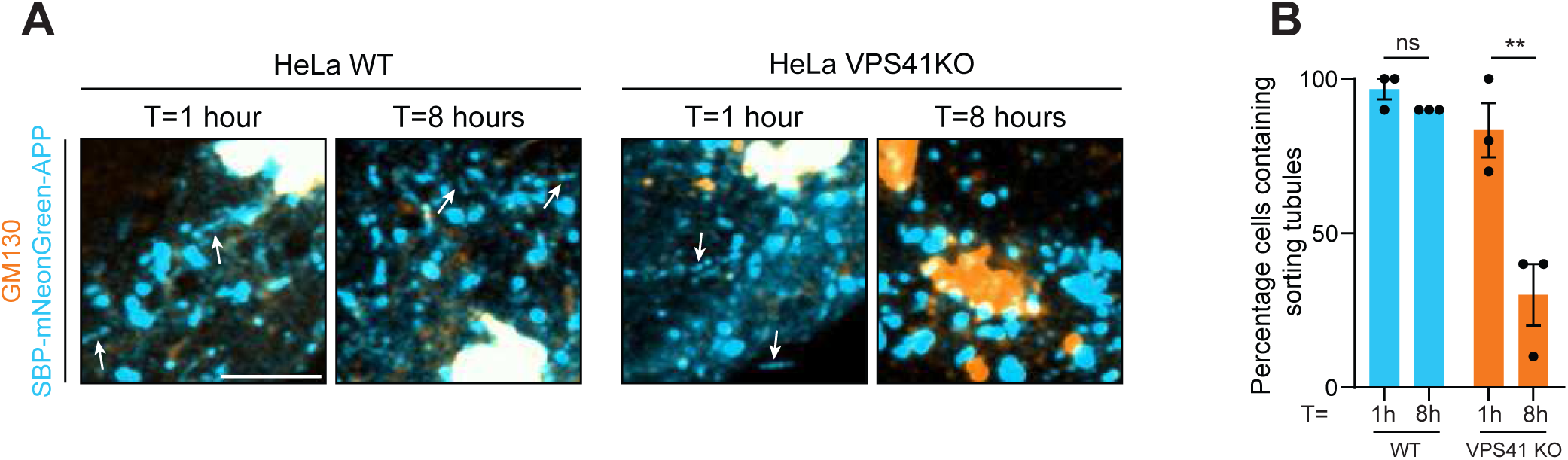
HOPS KO reduces formation of RUSH-APP recycling tubules. **A)** HeLa WT and VPS41 KO cells after 1h and 8h of RUSH-APP release from the ER. White arrows indicate APP positive tubules. **B)** Percentage of cells displaying APP-positive tubules at 1h and 8h of release. All scale bars indicate a region of 5 µm. Each dot represents the mean percentage of 10 cells (3 replicate experiments). Data is presented as mean values ± SEM. Ordinary two-way ANOVA was followed by Tukey’s correction for multiple comparisons was used in (B). ns = not significant, (P > 0.12), ** (P = 0.002).

**Figure S3.**
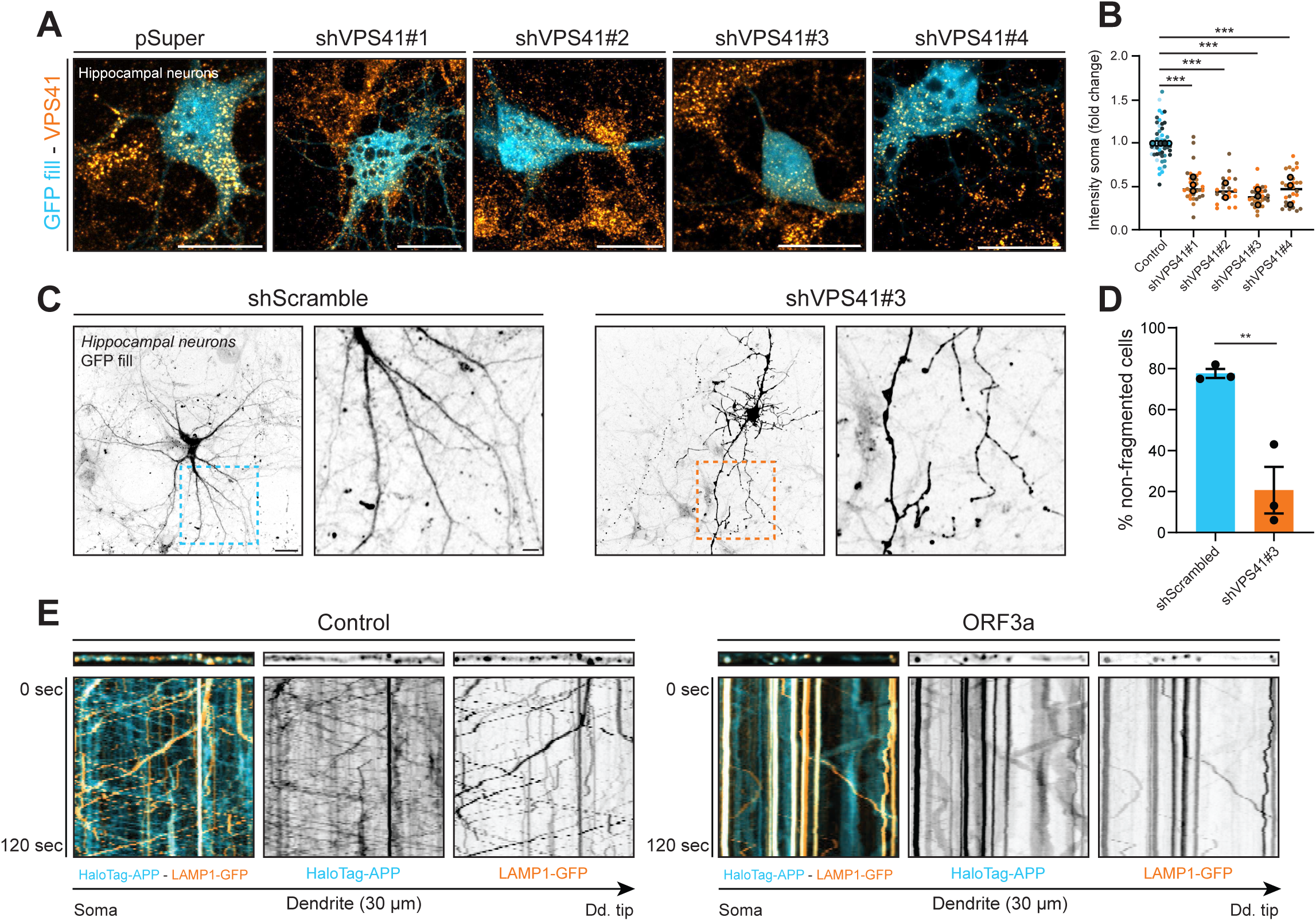
VPS41 knockdown induces neurite fragmentation and neuronal degeneration. **A)** Representative images of DIV8 hippocampal neurons transfected on DIV4 with GFP fill (blue) and pSuper / shScramble vector (control) or with different shRNAs targeting VPS41. Neurons were stained for endogenous VPS41 (orange). **B)** Quantification of VPS41 intensity in the soma of conditions in (A). **C)** Morphology of DIV15 hippocampal neurons transfected on DIV7 with GFP fill and shScramble or shVPS41 #3. Magnified regions (right). **D)** Quantification of the number of cells not showing neurite fragmentation after transfection with shScramble or shVPS41 as in (C). **E)** Kymographs of live cells expressing HaloTag-APP (N-terminal tag) and LAMP1-GFP in control and ORF3a expressing neurons. Dd. tip = dendrite tip. Dots in all dot plots indicate individual cells and lines indicate median value. Mean values of each replicate are indicated in dot blots by bold circles. Kruskal-Wallis test followed by Dunn’s correction for multiple comparisons was used in (B). Data is presented as mean values ± SEM in (D). Unpaired t test was conducted in (D). ** (P < 0.012), *** (P < 0.001).

**Figure S4.**
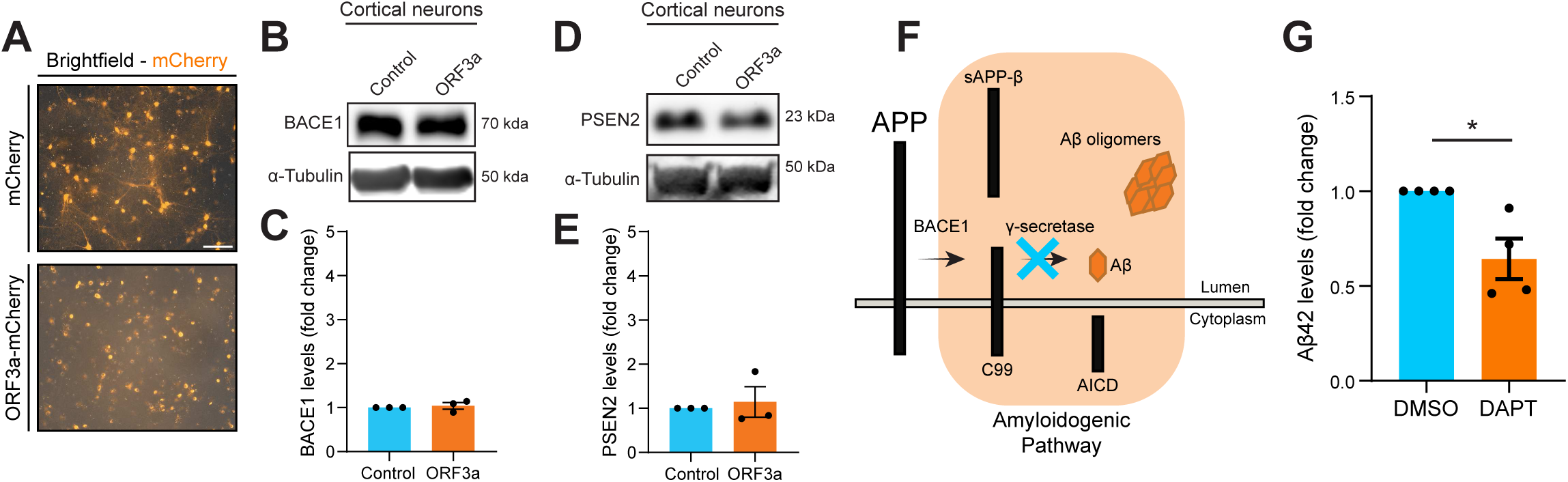
PSEN2 and BACE1 levels remain similar upon HOPS impairment. **A)** Representative widefield plus fluorescent images of DIV15 cortical neurons transduced on DIV7 using lentivirus for either mCherry (control) or ORF3a-mCherry (HOPS inhibition). **B-E)** Cortical neurons were transduced on DIV7 with lentivirus containing mCherry or ORF3a-mCherry and lysed on DIV15. Immunoblotting was performed for secretases BACE1 and PSEN2. Normalized protein levels are displayed in graphs below respective blots. Individual dots indicate replicates. **F)** Schematic of target of γ-secretase inhibitor DAPT. **G)** Measurement of Aβ42 levels in whole cell lysates from DIV15 neurons pre-treated for 16h with 25 µM γ-secretase inhibitor DAPT. Scale bar indicates a region of 100 µm. Unpaired t test with Welch’s correction was used in (G). Data is presented as mean values ± SEM in all graphs.

**Figure S5.**
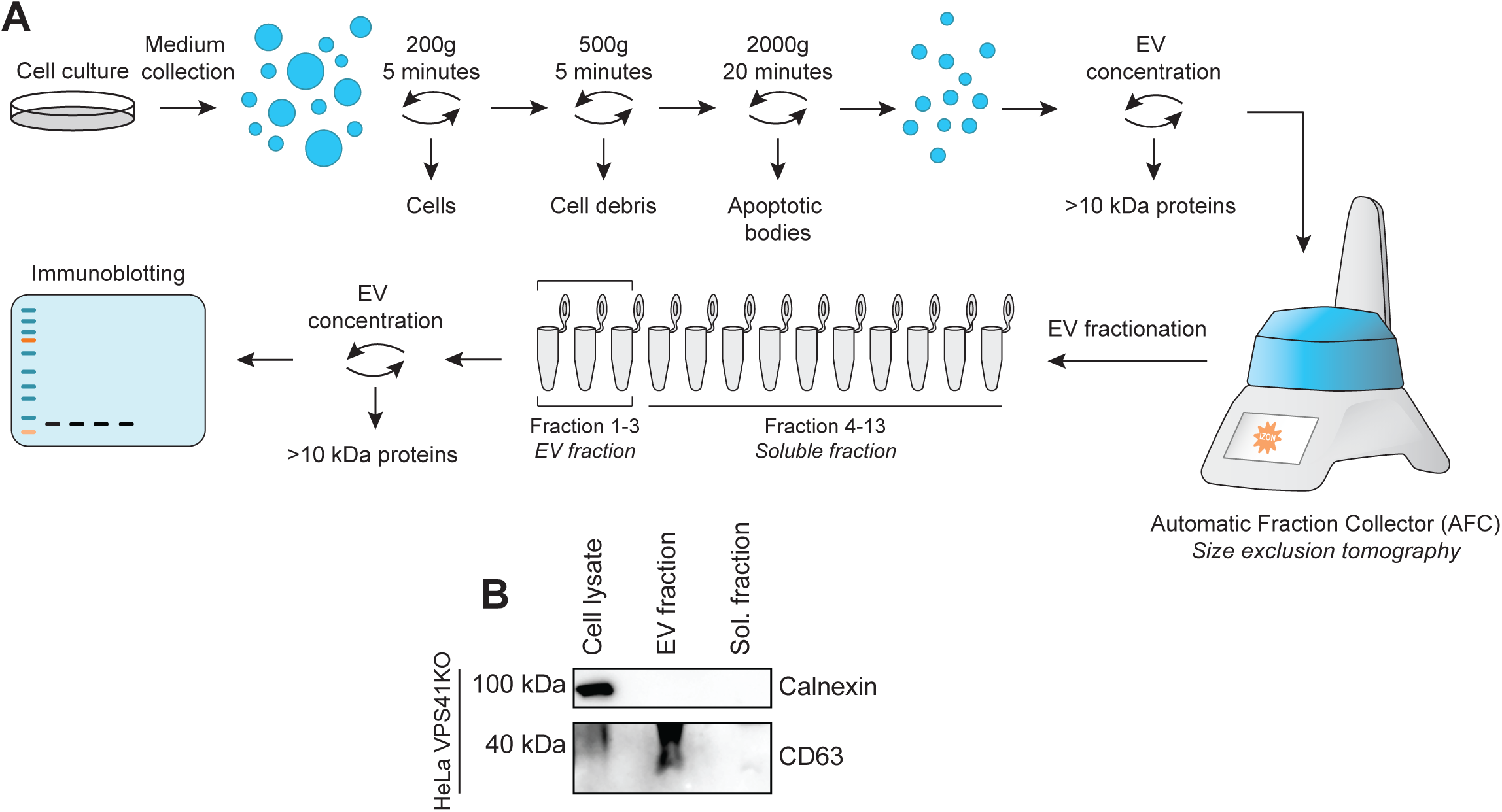
Validation of EV collection workflow. **A)** Schematic of isolation of EV and soluble fraction using automatic fraction collector (AFC), which is based on the size exclusion tomography principle. In all experiments, EVs were isolated from 25 mL of EV depleted culture medium after 72h of culture. Cell debris and apoptotic bodies were excluded from medium following indicated centrifugal steps. **B)** Proof of principle of EV isolation. ER marker (calnexin) is absent in only EV fraction and soluble fraction isolated from HeLa VPS41 KO cells. EV marker (CD63) is absent from soluble fraction isolated from HeLa VPS41 KO cells. CD63 can be detected as a smear due to glycosylation heterogeneity.

**Video S1. Release dynamics of RUSH-APP is similar in HeLa WT and VPS41 KO cells.** Representative videos (Spinning Disk microscope) of HeLa WT and Vps41 KO cells expressing RUSH-APP. Cells were imaged every minute for one hour. Biotin addition took place at T=0.

**Video S2. HOPS impairment in neurons increases C-terminal APP and LAMP1 colocalization in static LAMP1 compartments.** Representative videos (Spinning Disk microscope) of DIV8 hippocampal neurons and according cropped dendrites expressing LAMP1-GFP and APP-HaloTag (labeled using JFX650). Cells were imaged 2 minutes every second.

**Video S3. HOPS impairment in neurons increases N-terminal APP and LAMP1 colocalization in static LAMP1 compartments.** Representative videos (Spinning Disk microscope) of DIV8 hippocampal neurons and according cropped dendrites expressing LAMP1-GFP and HaloTag-APP (labeled using JFX650). Cells were imaged 2 minutes every second.

**Video S4. Secretion of APP fragments follow a CD63-positive route in HeLa VPS41 KO cells.** Representative videos (TIRF microscopy) of HeLa VPS41 KO cells expressing CD63-pHmScarlet and pHluorin-APP. Cells were imaged 3 minutes every 0.3 seconds.

## References

Van Acker, Z.P., M. Bretou, and W. Annaert. 2019. Endo-lysosomal dysregulations and late-onset Alzheimer’s disease: impact of genetic risk factors. Mol Neurodegener. 14:20. doi:10.1186/S13024-019-0323-7.

Allinquant, B., K.L. Moya, C. Bouillot, and A. Prochiantz. 1994. Amyloid precursor protein in cortical neurons: Coexistence of two pools differentially distributed in axons and dendrites and association with cytoskeleton. Journal of Neuroscience. 14:6842–6854. doi:10.1523/JNEUROSCI.14-11-06842.1994.

van der Beek, J., C. de Heus, P. Sanza, N. Liv, and J. Klumperman. 2024. Loss of the HOPS complex disrupts early-to-late endosome transition, impairs endosomal recycling and induces accumulation of amphisomes. Mol Biol Cell. 35:40–41. doi:10.1091/MBC.E23-08-0328/ASSET/IMAGES/LARGE/MBC-35-AR40-G010.JPEG.

van der Beek, J., C. Jonker, R. van der Welle, N. Liv, and J. Klumperman. 2019. CORVET, CHEVI and HOPS – Multisubunit tethers of the endo-lysosomal system in health and disease. J Cell Sci. 132. doi:10.1242/JCS.189134/57159.

van der Beek, J., and J. Klumperman. 2025. Trafficking to the lysosome: HOPS paves the way. Curr Opin Cell Biol. 94:102515. doi:10.1016/J.CEB.2025.102515.

Bentley, M., and G. Banker. 2016. The cellular mechanisms that maintain neuronal polarity. Nature Reviews Neuroscience 2016 17:10. 17:611–622. doi:10.1038/nrn.2016.100.

Bindels, D.S., L. Haarbosch, L. Van Weeren, M. Postma, K.E. Wiese, M. Mastop, S. Aumonier, G. Gotthard, A. Royant, M.A. Hink, and T.W.J. Gadella. 2016. MScarlet: A bright monomeric red fluorescent protein for cellular imaging. Nat Methods. 14:53–56. doi:10.1038/NMETH.4074.

Boncompain, G., S. Divoux, N. Gareil, H. De Forges, A. Lescure, L. Latreche, V. Mercanti, F. Jollivet, G. Raposo, and F. Perez. 2012. Synchronization of secretory protein traffic in populations of cells. Nature Methods 2012 9:5. 9:493–498. doi:10.1038/nmeth.1928.

Braulke, T., and J.S. Bonifacino. 2009. Sorting of lysosomal proteins. Biochimica et Biophysica Acta (BBA) - Molecular Cell Research. 1793:605–614. doi:10.1016/J.BBAMCR.2008.10.016.

Bretou, M., R. Sannerud, A. Escamilla-Ayala, T. Leroy, C. Vrancx, Z.P. Van Acker, A. Perdok, W. Vermeire, I. Vorsters, S. Van Keymolen, M. Maxson, B. Pavie, K. Wierda, E.L. Eskelinen, and W. Annaert. 2024. Accumulation of APP C-terminal fragments causes endolysosomal dysfunction through the dysregulation of late endosome to lysosome-ER contact sites. Dev Cell. 59:1571–1592.e9. doi:10.1016/J.DEVCEL.2024.03.030.

Britt, D.J., G.G. Farías, C.M. Guardia, and J.S. Bonifacino. 2016. Mechanisms of Polarized Organelle Distribution in Neurons. Front Cell Neurosci. 10:88. doi:10.3389/FNCEL.2016.00088.

Burd, C., and P.J. Cullen. 2014. Retromer: A Master Conductor of Endosome Sorting. Cold Spring Harb Perspect Biol. 6:a016774. doi:10.1101/CSHPERSPECT.A016774.

Burrinha, T., I. Martinsson, R. Gomes, A.P. Terrasso, G.K. Gouras, and C.G. Almeida. 2021. Upregulation of APP endocytosis by neuronal aging drives amyloid-dependent synapse loss. J Cell Sci. 134. doi:10.1242/JCS.255752/240244/AM/UP-REGULATION-OF-APP-ENDOCYTOSIS-BY-NEURONAL-AGING.

Butkovič, R., A.P. Walker, M.D. Healy, K.E. McNally, M. Liu, T. Veenendaal, K. Kato, N. Liv, J. Klumperman, B.M. Collins, and P.J. Cullen. 2024. Mechanism and regulation of cargo entry into the Commander endosomal recycling pathway. Nature Communications 2024 15:1. 15:1–15. doi:10.1038/s41467-024-50971-0.

Cataldo, A.M., C.M. Peterhoff, J.C. Troncoso, T. Gomez-Isla, B.T. Hyman, and R.A. Nixon. 2000. Endocytic Pathway Abnormalities Precede Amyloid β Deposition in Sporadic Alzheimer’s Disease and Down Syndrome: Differential Effects of APOE Genotype and Presenilin Mutations. Am J Pathol. 157:277. doi:10.1016/S0002-9440(10)64538-5.

Catsburg, L.A.E., M. Westra, A.M.L. van Schaik, and H.D. Macgillavry. 2022. Dynamics and nanoscale organization of the postsynaptic endocytic zone at excitatory synapses. Elife. 11:e74387. doi:10.7554/ELIFE.74387.

Chen, D., Q. Zheng, L. Sun, M. Ji, Y. Li, H. Deng, and H. Zhang. 2021. ORF3a of SARS-CoV-2 promotes lysosomal exocytosis-mediated viral egress. Dev Cell. 56:3250–3263.e5. doi:10.1016/J.DEVCEL.2021.10.006.

Chen, Y., D.C. Gershlick, S.Y. Park, and J.S. Bonifacino. 2017. Segregation in the Golgi complex precedes export of endolysosomal proteins in distinct transport carriers. J Cell Biol. 216:4141. doi:10.1083/JCB.201707172.

Chou, C.C., R. Vest, M.A. Prado, J. Wilson-Grady, J.A. Paulo, Y. Shibuya, P. Moran-Losada, T.T. Lee, J. Luo, S.P. Gygi, J.W. Kelly, D. Finley, M. Wernig, T. Wyss-Coray, and J. Frydman. 2025. Proteostasis and lysosomal repair deficits in transdifferentiated neurons of Alzheimer’s disease. Nature Cell Biology 2025 27:4. 27:619–632. doi:10.1038/s41556-025-01623-y.

Cone, A.S., S.N. Hurwitz, G.S. Lee, X. Yuan, Y. Zhou, Y. Li, and D.G. Meckes. 2020. Alix and Syntenin-1 direct amyloid precursor protein trafficking into extracellular vesicles. BMC Mol Cell Biol. 21:1–20. doi:10.1186/S12860-020-00302-0/FIGURES/9.

Currinn, H., B. Guscott, Z. Balklava, A. Rothnie, and T. Wassmer. 2016. APP controls the formation of PI(3,5)P2 vesicles through its binding of the PIKfyve complex. Cellular and Molecular Life Sciences. 73:393–408. doi:10.1007/S00018-015-1993-0.

Cuylen, S., C. Blaukopf, A.Z. Politi, T. Muller-Reichert, B. Neumann, I. Poser, J. Ellenberg, A.A. Hyman, and D.W. Gerlich. 2016. Ki-67 acts as a biological surfactant to disperse mitotic chromosomes. Nature. 535:308–312. doi:10.1038/NATURE18610.

Daly, J.L., C.M. Danson, P.A. Lewis, L. Zhao, S. Riccardo, L. Di Filippo, D. Cacchiarelli, D. Lee, S.J. Cross, K.J. Heesom, W.C. Xiong, A. Ballabio, J.R. Edgar, and P.J. Cullen. 2023. Multi-omic approach characterises the neuroprotective role of retromer in regulating lysosomal health. Nature Communications 2023 14:1. 14:1–19. doi:10.1038/s41467-023-38719-8.

Das, U., D.A. Scott, A. Ganguly, E.H. Koo, Y. Tang, and S. Roy. 2013. Activity-Induced Convergence of APP and BACE-1 in Acidic Microdomains via an Endocytosis-Dependent Pathway. Neuron. 79:447–460. doi:10.1016/J.NEURON.2013.05.035.

Das, U., L. Wang, A. Ganguly, J.M. Saikia, S.L. Wagner, E.H. Koo, and S. Roy. 2015. Visualizing APP and BACE-1 approximation in neurons yields insight into the amyloidogenic pathway. Nature Neuroscience 2016 19:1. 19:55–64. doi:10.1038/nn.4188.

Dharshini, S.A.P., Y.H. Taguchi, and M.M. Gromiha. 2019. Investigating the energy crisis in Alzheimer disease using transcriptome study. Scientific Reports 2019 9:1. 9:1–13. doi:10.1038/s41598-019-54782-y.

Droogers, W.J., J. Willems, H.D. Macgillavry, and A.P.H. de Jong. 2022. Duplex Labeling and Manipulation of Neuronal Proteins Using Sequential CRISPR/Cas9 Gene Editing. eNeuro. 9:ENEURO.0056-22.2022. doi:10.1523/ENEURO.0056-22.2022.

Edbauer, D., M. Willem, S. Lammich, H. Steiner, and C. Haass. 2002. Insulin-degrading Enzyme Rapidly Removes the β-Amyloid Precursor Protein Intracellular Domain (AICD). Journal of Biological Chemistry. 277:13389–13393. doi:10.1074/JBC.M111571200.

Edelstein, A.D., M.A. Tsuchida, N. Amodaj, H. Pinkard, R.D. Vale, and N. Stuurman. 2014. Advanced methods of microscope control using μManager software. Journal of Biological Methods 2014, 1(2), 1. 1:1. doi:10.14440/JBM.2014.36.

Ferguson, S.M. 2018. Axonal Transport and Maturation of Lysosomes. Curr Opin Neurobiol. 51:45. doi:10.1016/J.CONB.2018.02.020.

Fjorback, A.W., M. Seaman, C. Gustafsen, A. Mehmedbasic, S. Gokool, C. Wu, D. Militz, V. Schmidt, P. Madsen, J.R. Nyengaard, T.E. Willnow, E.I. Christensen, W.B. Mobley, A. Nykjær, and O.M. Andersen. 2012. Retromer Binds the FANSHY Sorting Motif in SorLA to Regulate Amyloid Precursor Protein Sorting and Processing. The Journal of Neuroscience. 32:1467. doi:10.1523/JNEUROSCI.2272-11.2012.

Gallwitz, L., F. Bleibaum, M. Voss, M. Schweizer, K. Spengler, D. Winter, F. Zöphel, S. Müller, S. Lichtenthaler, M. Damme, and P. Saftig. 2024. Cellular depletion of major cathepsin proteases reveals their concerted activities for lysosomal proteolysis. Cellular and Molecular Life Sciences. 81:1–21. doi:10.1007/S00018-024-05274-4/FIGURES/7.

García-Ayllón, M.S., I. Lopez-Font, C.P. Boix, J. Fortea, R. Sánchez-Valle, A. Lleó, J.L. Molinuevo, H. Zetterberg, K. Blennow, and J. Sáez-Valero. 2017. C-terminal fragments of the amyloid precursor protein in cerebrospinal fluid as potential biomarkers for Alzheimer disease. Scientific Reports 2017 7:1. 7:1–7. doi:10.1038/s41598-017-02841-7.

Goate, A., M.C. Chartier-Harlin, M. Mullan, J. Brown, F. Crawford, L. Fidani, L. Giuffra, A. Haynes, N. Irving, L. James, R. Mant, P. Newton, K. Rooke, P. Roques, C. Talbot, M. Pericak-Vance, A. Roses, R. Williamson, M. Rossor, M. Owen, and J. Hardy. 1991. Segregation of a missense mutation in the amyloid precursor protein gene with familial Alzheimer’s disease. Nature 1991 349:6311. 349:704–706. doi:10.1038/349704a0.

Gowrishankar, S., Y. Wu, and S.M. Ferguson. 2017. Impaired JIP3-dependent axonal lysosome transport promotes amyloid plaque pathology. Journal of Cell Biology. 216:3291–3305. doi:10.1083/JCB.201612148/VIDEO-2.

Gowrishankar, S., P. Yuan, Y. Wu, M. Schrag, S. Paradise, J. Grutzendler, P. De Camilli, and S.M. Ferguson. 2015. Massive accumulation of luminal protease-deficient axonal lysosomes at Alzheimer’s disease amyloid plaques. Proc Natl Acad Sci U S A. 112:E3699–E3708. doi:10.1073/PNAS.1510329112.

Griffin, E.F., X. Yan, K.A. Caldwell, and G.A. Caldwell. 2018. Distinct functional roles of Vps41-mediated neuroprotection in Alzheimer’s and Parkinson’s disease models of neurodegeneration. Hum Mol Genet. 27:4176–4193. doi:10.1093/HMG/DDY308.

Guo, Q., H. Li, S.S.K. Gaddam, N.J. Justice, C.S. Robertson, and H. Zheng. 2011. Amyloid Precursor Protein Revisited: NEURON-SPECIFIC EXPRESSION AND HIGHLY STABLE NATURE OF SOLUBLE DERIVATIVES. J Biol Chem. 287:2437. doi:10.1074/JBC.M111.315051.

Harrington, A.J., T.A. Yacoubian, S.R. Slone, K.A. Caldwell, and G.A. Caldwell. 2012. Functional Analysis of VPS41-Mediated Neuroprotection in Caenorhabditis elegans and Mammalian Models of Parkinson’s Disease. Journal of Neuroscience. 32:2142–2153. doi:10.1523/JNEUROSCI.2606-11.2012.

Hitschler, L., and T. Lang. 2022. The transmembrane domain of the amyloid precursor protein is required for antiamyloidogenic processing by α-secretase ADAM10. Journal of Biological Chemistry. 298. doi:10.1016/J.JBC.2022.101911/ATTACHMENT/53B9204C-1299-4D75-B369-C22CE7750561/MMC1.DOCX.

Hoe, H.S., H.K. Lee, and D.T.S. Pak. 2010. The Upside of APP at Synapses. CNS Neurosci Ther. 18:47. doi:10.1111/J.1755-5949.2010.00221.X.

Holstege, H., S.J. Van Der Lee, M. Hulsman, T.H. Wong, J.G.J. Van Rooij, M. Weiss, E. Louwersheimer, F.J. Wolters, N. Amin, A.G. Uitterlinden, A. Hofman, M.A. Ikram, J.C. Van Swieten, H. Meijers-Heijboer, W.M. Van Der Flier, M.J.T. Reinders, C.M. Van Duijn, and P. Scheltens. 2017. Characterization of pathogenic SORL1 genetic variants for association with Alzheimer’s disease: a clinical interpretation strategy. European Journal of Human Genetics 2017 25:8. 25:973–981. doi:10.1038/ejhg.2017.87.

Januário, Y.C., J. Eden, L.S. de Oliveira, R. De Pace, L.A. Tavares, M.E. da Silva-Januário, V.B. Apolloni, E.L. Wilby, R. Altmeyer, P. V. Burgos, S.A.L. Corrêa, D.C. Gershlick, and L.L.P. daSilva. 2022. Clathrin adaptor AP-1–mediated Golgi export of amyloid precursor protein is crucial for the production of neurotoxic amyloid fragments. Journal of Biological Chemistry. 298:102172. doi:10.1016/J.JBC.2022.102172.

Jensen, A.M.G., J. Raska, P. Fojtik, G. Monti, M. Lunding, S. Bartova, V. Pospisilova, S.J. van der Lee, J. Van Dongen, L. Bossaerts, C. Van Broeckhoven, O. Dols-Icardo, A. Lléo, S. Bellini, R. Ghidoni, M. Hulsman, G.A. Petsko, K. Sleegers, D. Bohaciakova, H. Holstege, and O.M. Andersen. 2024. The SORL1 p.Y1816C variant causes impaired endosomal dimerization and autosomal dominant Alzheimer’s disease. Proceedings of the National Academy of Sciences. 121:e2408262121. doi:10.1073/PNAS.2408262121/SUPPL_FILE/PNAS.2408262121.SAPP.PDF.

Jia, D., J.S. Zhang, F. Li, J. Wang, Z. Deng, M.A. White, D.G. Osborne, C. Phillips-Krawczak, T.S. Gomez, H. Li, A. Singla, E. Burstein, D.D. Billadeau, and M.K. Rosen. 2016. Structural and mechanistic insights into regulation of the retromer coat by TBC1d5. Nature Communications 2016 7:1. 7:1–11. doi:10.1038/ncomms13305.

Jiang, P., T. Nishimura, Y. Sakamaki, E. Itakura, T. Hatta, T. Natsume, and N. Mizushima. 2014. The HOPS complex mediates autophagosome–lysosome fusion through interaction with syntaxin 17. Mol Biol Cell. 25:1327. doi:10.1091/MBC.E13-08-0447.

Jonsson, T., J.K. Atwal, S. Steinberg, J. Snaedal, P. V. Jonsson, S. Bjornsson, H. Stefansson, P. Sulem, D. Gudbjartsson, J. Maloney, K. Hoyte, A. Gustafson, Y. Liu, Y. Lu, T. Bhangale, R.R. Graham, J. Huttenlocher, G. Bjornsdottir, O.A. Andreassen, E.G. Jonsson, A. Palotie, T.W. Behrens, O.T. Magnusson, A. Kong, U. Thorsteinsdottir, R.J. Watts, and K. Stefansson. 2012. A mutation in APP protects against Alzheimer’s disease and age-related cognitive decline. Nature 2012 488:7409. 488:96–99. doi:10.1038/nature11283.

van der Kant, R., and L.S.B. Goldstein. 2015. Cellular Functions of the Amyloid Precursor Protein from Development to Dementia. Dev Cell. 32:502–515. doi:10.1016/j.devcel.2015.01.022.

Kapitein, L.C., and C.C. Hoogenraad. 2011. Which way to go? Cytoskeletal organization and polarized transport in neurons. Molecular and Cellular Neuroscience. 46:9–20. doi:10.1016/J.MCN.2010.08.015.

Kapitein, L.C., M.A. Schlager, M. Kuijpers, P.S. Wulf, M. van Spronsen, F.C. MacKintosh, and C.C. Hoogenraad. 2010a. Mixed Microtubules Steer Dynein-Driven Cargo Transport into Dendrites. Current Biology. 20:290–299. doi:10.1016/J.CUB.2009.12.052.

Kapitein, L.C., M.A. Schlager, W.A. Van Der Zwan, P.S. Wulf, N. Keijzer, and C.C. Hoogenraad. 2010b. Probing intracellular motor protein activity using an inducible cargo trafficking assay. Biophys J. 99:2143–2152. doi:10.1016/j.bpj.2010.07.055.

Kaur, G., M. Pawlik, S.E. Gandy, M.E. Ehrlich, J.F. Smiley, and E. Levy. 2016. Lysosomal dysfunction in the brain of a mouse model with intraneuronal accumulation of carboxyl terminal fragments of the amyloid precursor protein. Molecular Psychiatry 2016 22:7. 22:981–989. doi:10.1038/mp.2016.189.

Klumperman, J. 2011. Architecture of the Mammalian Golgi. Cold Spring Harb Perspect Biol. 3:a005181. doi:10.1101/CSHPERSPECT.A005181.

Kwart, D., A. Gregg, C. Scheckel, E. Murphy, D. Paquet, M. Duffield, J. Fak, O. Olsen, R. Darnell, and M. Tessier-Lavigne. 2019. A Large Panel of Isogenic APP and PSEN1 Mutant Human iPSC Neurons Reveals Shared Endosomal Abnormalities Mediated by APP β-CTFs, Not Aβ. Neuron. 104:256–270.e5. doi:10.1016/J.NEURON.2019.07.010.

Laulagnier, K., C. Javalet, F.J. Hemming, M. Chivet, G. Lachenal, B. Blot, C. Chatellard, and R. Sadoul. 2017. Amyloid precursor protein products concentrate in a subset of exosomes specifically endocytosed by neurons. Cell Mol Life Sci. 75:757. doi:10.1007/S00018-017-2664-0.

Lauritzen, I., A. Bécot, A. Bourgeois, R. Pardossi-Piquard, M.G. Biferi, M. Barkats, and F. Checler. 2019. Targeting γ-secretase triggers the selective enrichment of oligomeric APP-CTFs in brain extracellular vesicles from Alzheimer cell and mouse models. Transl Neurodegener. 8:1–17. doi:10.1186/S40035-019-0176-6/FIGURES/7.

Lauritzen, I., R. Pardossi-Piquard, C. Bauer, E. Brigham, J.D. Abraham, S. Ranaldi, P. Fraser, P. St-George-Hyslop, O. Le Thuc, V. Espin, L. Chami, J. Dunys, and F. Checler. 2012. The β-Secretase-Derived C-Terminal Fragment of βAPP, C99, But Not Aβ, Is a Key Contributor to Early Intraneuronal Lesions in Triple-Transgenic Mouse Hippocampus. Journal of Neuroscience. 32:16243–16255. doi:10.1523/JNEUROSCI.2775-12.2012.

Li, C.H., N. Kersten, N. Özkan, D.T.M. Nguyen, M. Koppers, H. Post, M. Altelaar, and G.G. Farias. 2024. Spatiotemporal proteomics reveals the biosynthetic lysosomal membrane protein interactome in neurons. Nature Communications. 15:1–19. doi:10.1038/S41467-024-55052-W;TECHMETA=13,14,51,58,63,82;SUBJMETA=313,631,642,80;KWRD=MEMBRANE+TRAFFICKING,ORGANELLES.

Lois, C., E.J. Hong, S. Pease, E.J. Brown, and D. Baltimore. 2002. Germline transmission and tissue-specific expression of transgenes delivered by lentiviral vectors. Science (1979). 295:868–872. doi:10.1126/SCIENCE.1067081/SUPPL_FILE/1067081S9_THUMB.GIF.

Luo, M., J. Zhou, C. Sun, W. Chen, C. Fu, C. Si, Y. Zhang, Y. Geng, and Y. Chen. 2025. APP β-CTF triggers cell-autonomous synaptic toxicity independent of Aβ. Elife. 13. doi:10.7554/ELIFE.100968.2.

Maloney, J.A., T. Bainbridge, A. Gustafson, S. Zhang, R. Kyauk, P. Steiner, M. Van Der Brug, Y. Liu, J.A. Ernst, R.J. Watts, and J.K. Atwal. 2014. Molecular Mechanisms of Alzheimer Disease Protection by the A673T Allele of Amyloid Precursor Protein. Journal of Biological Chemistry. 289:30990–31000. doi:10.1074/JBC.M114.589069.

Mehmedbasic, A., S.K. Christensen, J. Nilsson, U. Rüetschi, C. Gustafsen, A.S.A. Poulsen, R.W. Rasmussen, A.N. Fjorback, G. Larson, and O.M. Andersen. 2015. SorLA complement-type repeat domains protect the amyloid precursor protein against processing. J Biol Chem. 290:3359–3376. doi:10.1074/JBC.M114.619940.

Miao, G., H. Zhao, Y. Li, M. Ji, Y. Chen, Y. Shi, Y. Bi, P. Wang, and H. Zhang. 2020. ORF3a of the COVID-19 virus SARS-CoV-2 blocks HOPS complex-mediated assembly of the SNARE complex required for autolysosome formation. Dev Cell. 56:427. doi:10.1016/J.DEVCEL.2020.12.010.

Miserey-Lenkei, S., K. Trajkovic, J.M. D’Ambrosio, A.J. Patel, A. Čopič, P. Mathur, K. Schauer, B. Goud, V. Albanèse, R. Gautier, M. Subra, D. Kovacs, H. Barelli, and B. Antonny. 2021. A comprehensive library of fluorescent constructs of SARS-CoV-2 proteins and their initial characterisation in different cell types. Biol Cell. 113:311. doi:10.1111/BOC.202000158.

Monfrini, E., M. Zech, D. Steel, M.A. Kurian, J. Winkelmann, and A. Di Fonzo. 2021. HOPS-associated neurological disorders (HOPSANDs): linking endolysosomal dysfunction to the pathogenesis of dystonia. Brain. 144:2610–2615. doi:10.1093/BRAIN/AWAB161.

Mueller-Steiner, S., Y. Zhou, H. Arai, E.D. Roberson, B. Sun, J. Chen, X. Wang, G. Yu, L. Esposito, L. Mucke, and L. Gan. 2006. Antiamyloidogenic and Neuroprotective Functions of Cathepsin B: Implications for Alzheimer’s Disease. Neuron. 51:703–714. doi:10.1016/J.NEURON.2006.07.027/ATTACHMENT/9FB470D7-7551-49EA-8481-8EABA2B14C7B/MMC1.PDF.

Müller, U.C., T. Deller, and M. Korte. 2017. Not just amyloid: physiological functions of the amyloid precursor protein family. Nature Reviews Neuroscience 2017 18:5. 18:281–298. doi:10.1038/nrn.2017.29.

Park, H., F. V. Hundley, Q. Yu, K.A. Overmyer, D.R. Brademan, L. Serrano, J.A. Paulo, J.C. Paoli, S. Swarup, J.J. Coon, S.P. Gygi, and J. Wade Harper. 2022. Spatial snapshots of amyloid precursor protein intramembrane processing via early endosome proteomics. Nature Communications 2022 13:1. 13:1–21. doi:10.1038/s41467-022-33881-x.

Perez-Gonzalez, R., S.A. Gauthier, A. Kumar, and E. Levy. 2012. The Exosome Secretory Pathway Transports Amyloid Precursor Protein Carboxyl-terminal Fragments from the Cell into the Brain Extracellular Space. Journal of Biological Chemistry. 287:43108–43115. doi:10.1074/JBC.M112.404467.

Pietilä, M., P. Sahgal, E. Peuhu, N.Z. Jäntti, I. Paatero, E. Närvä, H. Al-Akhrass, J. Lilja, M. Georgiadou, O.M. Andersen, A. Padzik, H. Sihto, H. Joensuu, M. Blomqvist, I. Saarinen, P.J. Boström, P. Taimen, and J. Ivaska. 2019. SORLA regulates endosomal trafficking and oncogenic fitness of HER2. Nature Communications 2019 10:1. 10:1–16. doi:10.1038/s41467-019-10275-0.

Polanco, J.C., Y. Akimov, A. Fernandes, A. Briner, G.R. Hand, M. van Roijen, G. Balistreri, and J. Götz. 2022. CRISPRi screening reveals regulators of tau pathology shared between exosomal and vesicle-free tau. Life Sci Alliance. 6:e202201689. doi:10.26508/LSA.202201689.

Pols, M.S., E. Van Meel, V. Oorschot, C. Ten Brink, M. Fukuda, M.G. Swetha, S. Mayor, and J. Klumperman. 2013. hVps41 and VAMP7 function in direct TGN to late endosome transport of lysosomal membrane proteins. Nature Communications 2013 4:1. 4:1–12. doi:10.1038/ncomms2360.

Raghavan, N.S., A.M. Brickman, H. Andrews, J.J. Manly, N. Schupf, R. Lantigua, C.J. Wolock, S. Kamalakaran, S. Petrovski, G. Tosto, B.N. Vardarajan, D.B. Goldstein, and R. Mayeux. 2018. Whole-exome sequencing in 20,197 persons for rare variants in Alzheimer’s disease. Ann Clin Transl Neurol. 5:832–842. doi:10.1002/ACN3.582.

Rink, J., E. Ghigo, Y. Kalaidzidis, and M. Zerial. 2005. Rab conversion as a mechanism of progression from early to late endosomes. Cell. 122:735–749. doi:10.1016/J.CELL.2005.06.043.

Rogaev, E.I., R. Sherrington, E.A. Rogaeva, G. Levesque, M. Ikeda, Y. Liang, H. Chi, C. Lin, K. Holman, T. Tsuda, L. Mar, S. Sorbi, B. Nacmias, S. Piacentini, L. Amaducci, I. Chumakov, D. Cohen, L. Lannfelt, P.E. Fraser, J.M. Rommens, and P.H.S. George-Hyslop. 1995. Familial Alzheimer’s disease in kindreds with missense mutations in a gene on chromosome 1 related to the Alzheimer’s disease type 3 gene. Nature 1995 376:6543. 376:775–778. doi:10.1038/376775a0.

Rojas, R., T. Van Vlijmen, G.A. Mardones, Y. Prabhu, A.L. Rojas, S. Mohammed, A.J.R. Heck, G. Raposo, P. Van Der Sluijs, and J.S. Bonifacino. 2008. Regulation of retromer recruitment to endosomes by sequential action of Rab5 and Rab7. J Cell Biol. 183:513. doi:10.1083/JCB.200804048.

Salvadores, N., C. Gerónimo-Olvera, and F.A. Court. 2020. Axonal Degeneration in AD: The Contribution of Aβ and Tau. Front Aging Neurosci. 12:581767. doi:10.3389/FNAGI.2020.581767.

Sannerud, R., I. Declerck, A. Peric, T. Raemaekers, G. Menendez, L. Zhou, B. Veerle, K. Coen, S. Munck, B. De Strooper, G. Schiavo, and W. Annaert. 2011. ADP ribosylation factor 6 (ARF6) controls amyloid precursor protein (APP) processing by mediating the endosomal sorting of BACE1. Proc Natl Acad Sci U S A. 108:E559–E568. doi:10.1073/PNAS.1100745108/SUPPL_FILE/PNAS.201100745SI.PDF.

Sannerud, R., C. Esselens, P. Ejsmont, R. Mattera, L. Rochin, A.K. Tharkeshwar, G. De Baets, V. De Wever, R. Habets, V. Baert, W. Vermeire, C. Michiels, A.J. Groot, R. Wouters, K. Dillen, K. Vints, P. Baatsen, S. Munck, R. Derua, E. Waelkens, G.S. Basi, M. Mercken, M. Vooijs, M. Bollen, J. Schymkowitz, F. Rousseau, J.S. Bonifacino, G. Van Niel, B. De Strooper, and W. Annaert. 2016. Restricted Location of PSEN2/γ-Secretase Determines Substrate Specificity and Generates an Intracellular Aβ Pool. Cell. 166:193–208. doi:10.1016/J.CELL.2016.05.020.

Sanzà, P., J. van der Beek, D. Draper, C. de Heus, T. Veenendaal, C. Ten Brink, G.G. Farías, N. Liv, and J. Klumperman. 2025. VPS41 recruits biosynthetic LAMP-positive vesicles through interaction with Arl8b. J Cell Biol. 224. doi:10.1083/JCB.202405002/277255.

Sardar Sinha, M., A. Ansell-Schultz, L. Civitelli, C. Hildesjö, M. Larsson, L. Lannfelt, M. Ingelsson, and M. Hallbeck. 2018. Alzheimer’s disease pathology propagation by exosomes containing toxic amyloid-beta oligomers. Acta Neuropathol. 136:41–56. doi:10.1007/S00401-018-1868-1/FIGURES/2.

Schleinitz, A., L.A. Pöttgen, T. Keren-Kaplan, J. Pu, P. Saftig, J.S. Bonifacino, A. Haas, and A. Jeschke. 2023. Consecutive functions of small GTPases guide HOPS-mediated tethering of late endosomes and lysosomes. Cell Rep. 42. doi:10.1016/J.CELREP.2022.111969/ATTACHMENT/835269E9-79A5-4EFA-97A1-7AC8014F33F9/MMC2.PDF.

Shelke, G.V., C.D. Williamson, M. Jarnik, and J.S. Bonifacino. 2023. Inhibition of endolysosome fusion increases exosome secretion. J Cell Biol. 222. doi:10.1083/JCB.202209084.

Sherrington, R., E.I. Rogaev, Y. Liang, E.A. Rogaeva, G. Levesque, M. Ikeda, H. Chi, C. Lin, G. Li, K. Holman, T. Tsuda, L. Mar, J.F. Foncin, A.C. Bruni, M.P. Montesi, S. Sorbi, I. Rainero, L. Pinessi, L. Nee, I. Chumakov, D. Pollen, A. Brookes, P. Sanseau, R.J. Polinsky, W. Wasco, H.A.R. Da Silva, J.L. Haines, M.A. Pericak-Vance, R.E. Tanzi, A.D. Roses, P.E. Fraser, J.M. Rommens, and P.H. St George-Hyslop. 1995. Cloning of a gene bearing missense mutations in early-onset familial Alzheimer’s disease. Nature 1995 375:6534. 375:754–760. doi:10.1038/375754a0.

Slot, J.W., and H.J. Geuze. 2007. Cryosectioning and immunolabeling. Nature Protocols 2007 2:10. 2:2480–2491. doi:10.1038/nprot.2007.365.

Small, S.A., K. Kent, A. Pierce, C. Leung, M.S. Kang, H. Okada, L. Honig, J.P. Vonsattel, and T.W. Kim. 2005. Model-guided microarray implicates the retromer complex in Alzheimer’s disease. Ann Neurol. 58:909–919. doi:10.1002/ANA.20667.

Small, S.A., S. Simoes-Spassov, R. Mayeux, and G.A. Petsko. 2017. Endosomal traffic jams represent a pathogenic hub and therapeutic target in Alzheimer’s disease. Trends Neurosci. 40:592. doi:10.1016/J.TINS.2017.08.003.

Steel, D., M. Zech, C. Zhao, K.E.S. Barwick, D. Burke, D. Demailly, K.R. Kumar, G. Zorzi, N. Nardocci, R. Kaiyrzhanov, M. Wagner, A. Iuso, R. Berutti, M. Škorvánek, J. Necpál, R. Davis, S. Wiethoff, K. Mankad, S. Sudhakar, A. Ferrini, S. Sharma, E.J. Kamsteeg, M.A. Tijssen, C. Verschuuren, M.E. van Egmond, J.M. Flowers, M. McEntagart, A. Tucci, P. Coubes, B.I. Bustos, P. Gonzalez-Latapi, S. Tisch, P. Darveniza, K.M. Gorman, K.J. Peall, K. Bötzel, J.C. Koch, T. Kmieć, B. Plecko, S. Boesch, B. Haslinger, R. Jech, B. Garavaglia, N. Wood, H. Houlden, P. Gissen, S.J. Lubbe, C.M. Sue, L. Cif, N.E. Mencacci, G. Anderson, M.A. Kurian, and J. Winkelmann. 2020. Loss-of-Function Variants in HOPS Complex Genes VPS16 and VPS41 Cause Early Onset Dystonia Associated with Lysosomal Abnormalities. Ann Neurol. 88:867–877. doi:10.1002/ANA.25879.

Subramanian, J., J.C. Savage, and M.È. Tremblay. 2020. Synaptic Loss in Alzheimer’s Disease: Mechanistic Insights Provided by Two-Photon in vivo Imaging of Transgenic Mouse Models. Front Cell Neurosci. 14:592607. doi:10.3389/FNCEL.2020.592607.

Swetha, M.G., V. Sriram, K.S. Krishnan, V.M. J Oorschot, C. ten Brink, J. Klumperman, and S. Mayor. 2011. Lysosomal Membrane Protein Composition, Acidic pH and Sterol Content are Regulated via a Light-Dependent Pathway in Metazoan Cells. Traffic. 12:1037–1055. doi:10.1111/j.1600-0854.2011.01214.x.

Tan, J.Z.A., L. Fourriere, J. Wang, F. Perez, G. Boncompain, and P.A. Gleesona. 2020. Distinct anterograde trafficking pathways of BACE1 and amyloid precursor protein from the TGN and the regulation of amyloid-β production. Mol Biol Cell. 31:27–44. doi:10.1091/MBC.E19-09-0487.

Tan, J.Z.A., and P.A. Gleeson. 2019a. The trans-Golgi network is a major site for α-secretase processing of amyloid precursor protein in primary neurons. Journal of Biological Chemistry. 294:1618–1631. doi:10.1074/JBC.RA118.005222.

Tan, J.Z.A., and P.A. Gleeson. 2019b. The role of membrane trafficking in the processing of amyloid precursor protein and production of amyloid peptides in Alzheimer’s disease. Biochimica et Biophysica Acta (BBA) - Biomembranes. 1861:697–712. doi:10.1016/J.BBAMEM.2018.11.013.

Toh, W.H., P.Z. Cheryl Chia, M. Iqbal Hossain, and P.A. Gleeson. 2018. GGA1 regulates signal-dependent sorting of BACE1 to recycling endosomes, which moderates A? production. Mol Biol Cell. 29:191–208. doi:10.1091/MBC.E17-05-0270/ASSET/IMAGES/LARGE/MBC-29-191-G010.JPEG.

Toh, W.H., J.Z.A. Tan, K.L. Zulkefli, F.J. Houghton, and P.A. Gleeson. 2017. Amyloid precursor protein traffics from the Golgi directly to early endosomes in an Arl5b- and AP4-dependent pathway. Traffic. 18:159–175. doi:10.1111/TRA.12465.

Vassar, R., B.D. Bennett, S. Babu-Khan, S. Kahn, E.A. Mendiaz, P. Denis, D.B. Teplow, S. Ross, P. Amarante, R. Loeloff, Y. Luo, S. Fisher, J. Fuller, S. Edenson, J. Lile, M.A. Jarosinski, A.L. Biere, E. Curran, T. Burgess, J.C. Louis, F. Collins, J. Treanor, G. Rogers, and M. Citron. 1999. β-Secretase cleavage of Alzheimer’s amyloid precursor protein by the transmembrane aspartic protease BACE. Science (1979). 286:735–741. doi:10.1126/SCIENCE.286.5440.735/ASSET/FF706FE5-60AA-4375-902A-FE30A5CFF516/ASSETS/GRAPHIC/SE4297936008.JPEG.

Verweij, F.J., M.P. Bebelman, C.R. Jimenez, J.J. Garcia-Vallejo, H. Janssen, J. Neefjes, J.C. Knol, R. de Goeij-de Haas, S.R. Piersma, S.R. Baglio, M. Verhage, J.M. Middeldorp, A. Zomer, J. van Rheenen, M.G. Coppolino, I. Hurbain, G. Raposo, M.J. Smit, R.F.G. Toonen, G. van Niel, and D.M. Pegtel. 2018. Quantifying exosome secretion from single cells reveals a modulatory role for GPCR signaling. Journal of Cell Biology. 217:1129–1142. doi:10.1083/JCB.201703206/VIDEO-9.

Vieira, S.I., S. Rebelo, H. Esselmann, J. Wiltfang, J. Lah, R. Lane, S.A. Small, S. Gandy, E.F. Da Cruz E Silva, and O.A.B. Da Cruz E Silva. 2010. Retrieval of the Alzheimer’s amyloid precursor protein from the endosome to the TGN is S655 phosphorylation state-dependent and retromer-mediated. Mol Neurodegener. 5:40. doi:10.1186/1750-1326-5-40.

Villegas, C., V. Muresan, and Z. Ladescu Muresan. 2013. Dual-tagged amyloid-β precursor protein reveals distinct transport pathways of its N- and C-terminal fragments. Hum Mol Genet. 23:1631. doi:10.1093/HMG/DDT555.

Walia, K., A. Sharma, S. Paul, P. Chouhan, G. Kumar, R. Ringe, M. Sharma, and A. Tuli. 2024. SARS-CoV-2 virulence factor ORF3a blocks lysosome function by modulating TBC1D5-dependent Rab7 GTPase cycle. Nat Commun. 15. doi:10.1038/S41467-024-46417-2.

van der Welle, R.E.N., R. Jobling, C. Burns, P. Sanza, J.A. van der Beek, A. Fasano, L. Chen, F.J. Zwartkruis, S. Zwakenberg, E.F. Griffin, C. ten Brink, T. Veenendaal, N. Liv, C.M.A. van Ravenswaaij-Arts, H.H. Lemmink, R. Pfundt, S. Blaser, C. Sepulveda, A.M. Lozano, G. Yoon, T. Santiago-Sim, C.S. Asensio, G.A. Caldwell, K.A. Caldwell, D. Chitayat, and J. Klumperman. 2021. Neurodegenerative VPS41 variants inhibit HOPS function and mTORC1-dependent TFEB/TFE3 regulation. EMBO Mol Med. 13. doi:10.15252/EMMM.202013258.

Wilson, D.M., M.R. Cookson, L. Van Den Bosch, H. Zetterberg, D.M. Holtzman, and I. Dewachter. 2023. Hallmarks of neurodegenerative diseases. Cell. 186:693–714. doi:10.1016/J.CELL.2022.12.032.

Xiao, T., W. Zhang, B. Jiao, C.Z. Pan, X. Liu, and L. Shen. 2017. The role of exosomes in the pathogenesis of Alzheimer’ disease. Transl Neurodegener. 6:1–6. doi:10.1186/S40035-017-0072-X/METRICS.

Xie, J., E.N. Heim, M. Crite, and D. DiMaio. 2020. TBC1D5-Catalyzed Cycling of Rab7 Is Required for Retromer-Mediated Human Papillomavirus Trafficking during Virus Entry. Cell Rep. 31:107750. doi:10.1016/J.CELREP.2020.107750/ATTACHMENT/69B6D937-38C3-4B6E-98FA-60ACEE5D0D7B/MMC2.PDF.

Yap, C.C., L. Digilio, L.P. McMahon, A.D.R. Garcia, and B. Winckler. 2018. Degradation of dendritic cargos requires Rab7-dependent transport to somatic lysosomes. Journal of Cell Biology. 217:3141–3159. doi:10.1083/JCB.201711039.

Yap, C.C., A.J. Mason, and B. Winckler. 2022. “Dynamics and distribution of endosomes and lysosomes in dendrites.” Curr Opin Neurobiol. 74:102537. doi:10.1016/J.CONB.2022.102537.

Zhang, H., T. Huang, Y. Hong, W. Yang, X. Zhang, H. Luo, H. Xu, and X. Wang. 2018. The Retromer Complex and Sorting Nexins in Neurodegenerative Diseases. Front Aging Neurosci. 10:79. doi:10.3389/FNAGI.2018.00079.

Zhang, J., A. Ejikemeuwa, V. Gerzanich, M. Nasr, Q. Tang, J.M. Simard, and R.Y. Zhao. 2022. Understanding the Role of SARS-CoV-2 ORF3a in Viral Pathogenesis and COVID-19. Front Microbiol. 13:854567. doi:10.3389/FMICB.2022.854567/XML.

Zhang, J., V. Lachance, A. Schaffner, X. Li, A. Fedick, L.E. Kaye, J. Liao, J. Rosenfeld, N. Yachelevich, M.L. Chu, W.G. Mitchell, R.G. Boles, E. Moran, M. Tokita, E. Gorman, K. Bagley, W. Zhang, F. Xia, M. Leduc, Y. Yang, C. Eng, L.J. Wong, R. Schiffmann, G.A. Diaz, R. Kornreich, R. Thummel, M. Wasserstein, Z. Yue, and L. Edelmann. 2016. A Founder Mutation in VPS11 Causes an Autosomal Recessive Leukoencephalopathy Linked to Autophagic Defects. PLoS Genet. 12:e1005848. doi:10.1371/JOURNAL.PGEN.1005848.

